# Insight into Malt1 activation mechanism through synergetic approach of AlphaFold, MD Simulation and NMR dynamic analyses

**DOI:** 10.64898/2025.12.17.694852

**Authors:** Dmitry Lesovoy, Tatiana Agback, Konstantin Roshchin, Tatyana Sandalova, Adnane Achour, Xiao Han, Alexander Lomzov, Vladislav Orekhov, Peter Agback

## Abstract

Mucosa-associated lymphoid tissue lymphoma translocation protein 1 (MALT1) is a central regulator of immune signalling, yet the conformational dynamics governing its activation remain poorly defined. Building on our earlier solution-state analysis of apo MALT1(PCASP-Ig3)_339–719_, which revealed domain flexibility, dynamic autoinhibition, and sensitivity to physiological ionic conditions, we combine NMR relaxation, molecular dynamics simulations, and ensemble modelling to delineate how solution environment reshapes its conformational landscape. Because most structural information derives from dimeric or inhibitor-bound states, the behaviour of monomeric, ligand-free MALT1 in physiological solution has remained unclear. Here, MD simulations show that low-salt conditions drive all trajectories toward a unified inactive-like ensemble, marked by inward rotation of W580 and coordinated rearrangements of Loop 2 and Loop 3, indicating that the inactive state is energetically favoured and its reactivation kinetically suppressed. Physiological ionic strength partially restores access to active-like loop motions, aligning with NMR evidence that sodium modulates catalytic readiness. In contrast, high-salt conditions rigidify the PCASP–Ig3 module, suppressing loop fluctuations and preventing active–inactive transitions, thereby strongly enriching the active-state population.

Importantly, the combined MD–NMR analysis demonstrates that the NMR-initiated ensembles provide the most faithful representation of backbone and loop dynamics under low-salt conditions, capturing substrate-independent loop rearrangements, stable hydrophobic-core behaviour, and the intrinsic transitions that shape MALT1’s conformational equilibrium.

Together, these findings identify ionic strength as a key regulator of MALT1 conformational equilibria,, highlighting how loop dynamics and domain flexibility tune its proteolytic competence and providing a dynamic framework for future structure-based modulation of MALT1 activity.

## 1.0 Introduction

Mucosa-associated lymphoid tissue lymphoma-translocation protein 1 (MALT1), a crucial human paracaspase forms a complex with CARD11 and BCL10 (CMB)^1–3^ that plays a central role in activating the NF-κB signalling pathway following antigen receptor stimulation. It is essential for the survival, proliferation, and function of B and T cells, making it a promising therapeutic target for lymphoma, cancer, and autoimmune diseases^4–11^. CBM complex assembly activates cysteine dependent endopeptidase activity of MALT1, which modulates immune responses by specific cleavage of several proteins involved in cell signaling, adhesion, transcription and mRNA regulation^3^.

MALT1 has a multi-domain structure consisting of an N-terminal death domain (DD), two immunoglobulin-like domains (Ig1, Ig2), a paracaspase (PCASP) domain^12, 13^, which contains the catalytic dyad H415/C464^14, 15^, third immunoglobulin-like Ig3 domain and about 100 C-terminal residues with unclear domain structure. Although all of these domains contribute to MALT1 function in cells, a two-domain construct containing only the paracaspase and Ig3 domains shows in vitro peptidase activity at high concentration of sodium citrate comparable with full-length protein^13^.

Over the past two decades, sustained research in molecular biology and biophysical analysis has significantly advanced our understanding of MALT1 activation mechanisms (see most recent reviews^3, 8, 9, 16–18^. Initially identified as a distant caspase homolog^19, 20^ MALT1 was thought to follow the two-step activation model: first, dimerization of the paracapase domain, followed by release from Ig3-mediated autoinhibition upon substrate binding^21^.

Several lines of evidence support this model. Earlier X-ray crystallographic studies of a truncated MALT1 construct containing the PCASP and Ig3 domains, MALT1(PCASP-Ig3)₃₃₉–₇₁₉, revealed an auto inhibited, monomeric state^13^ with dimer formation inducible by peptide ligands^12, 13^. LS-MS and chromatographic analyses confirmed the monomeric nature of this construct during purification. Nevertheless, both biochemical and crystallographic studies demonstrated that monomeric MALT1 can spontaneously form dimers in solution^22^, reinforcing the idea that dimerization is essential for activation.

These findings indicate that activation involves a conformational transition driven by regulatory interactions and ligand binding, which together stabilize the active form and enhance catalytic function^12, 13^.

Currently available X-ray crystal structures of MALT1, either in its active form bound to predominantly to peptide-based inhibitors^12, 13, 23, 24^ or in its inactive form bound to allosteric ligands^24–28^, provide structural snapshots that reveal significant conformational changes in the L2, L3, and L4 loops upon ligand binding.

Enzymes do not toggle between defined single inactive and active states, they are dynamic systems and exist as ensembles of various conformations. However, until recently, the conformational flexibility of MALT1 and interplay between its structural domains have received limited attention. The role of protein dynamics in regulating MALT1 activation remains poorly understood and has not been a primary focus of earlier studies.

Loops play very often the central roles in the enzyme catalysis. As flexible connectors between secondary-structure elements, they strongly influence catalysis, substrate recognition, allosteric regulation, and the global dynamical behaviour of enzyme scaffolds^29–37^. Their frequent evolutionary conservation underscores their functional importance^38^.

Our previous work on near-complete assignments of the resonances in NMR spectra of the apo form of human MALT1, (PCASP-Ig3)₃₃₉–₇₁₉^39^, and subsequently extended to IVL-methyl side-chain assignments for both wild-type and E549A mutant variants (Biological Magnetic Resonance Data Bank accession code BMRB 52265 (http://www.bmrb.wisc.edu/)^40^ laid the strong base for probing the protein internal dynamics using experimental relaxation data (R₁, R₂, and NOE) and thereby for validating conformational ensembles obtained from MD simulations. The strong base for probing the protein’s internal dynamics using experimental relaxation data

Based on these data, our recent NMR studies^41^ demonstrated that the MALT1(PCASP-Ig3)₃₃₉–₇₁₉ construct behaves as a monomer under physiological salt conditions, and that its PCASP and Ig3 domains show semi-independent motion that contrasts with the compact interface observed in the crystal structures^13^.

Complementary AlphaFold2 and AlphaFold3 modelling of the apo form of MALT1(PCASP-Ig3)₃₃₉–₇₁₉ predicted a set of conformational families based on activity of PCASP domain and the orientation of W580. Although AlphaFold lacks environmental context, its predictions likely reflect evolutionary and structural biases from the Protein Data Bank. Notably, according to crystallographic observations, the active state of the PCASP domain is observed in ligand-bound forms at the active site, highlighting the potential for functional conformations to exist in solution and to be influenced by subtle environmental changes, such as the presence of kosmotropic salts.^14^. This underscores the importance of recognizing the potential for shifts in populations within the conformational ensemble in solution under different environmental conditions.

The influence of physiological electrolytes, such as sodium, on MALT1(PCASP-Ig3)₃₃₉–₇₁₉ mediated activation remains unclear. However, conducting NMR experiments at elevated salt concentrations is technically challenging due to the construct’s relatively large size, increased spectral complexity, and compromised relaxation properties. In contrast, molecular dynamics (MD) simulations provide a complementary approach by enabling the investigation of MALT1 behaviour under high-salt conditions. These simulations offer valuable insights into the interplay of electrostatic and hydrophobic interactions, which are otherwise difficult to resolve experimentally in this context.

The present study has the following objectives: use the MD trajectories (i) to determine the conformation ensemble dominated intramolecular dynamical contributions to spin relaxation of backbone and of side-chain methyl groups of MALT1(PCASP-Ig3)₃₃₉–₇₁₉ at physiological condition, low salt concentration; (ii) to evaluate the possibility of transition between inactive and active conformations of protein, (iii) establish the influence on the conformation state of MALT1 environment conditions such as low and high salt concentrations.

Herein, comparisons between MD simulations and NMR spin relaxation measurements were performed using backbone NH and side-chain methyl relaxation parameters for MALT1(PCASP-Ig3)₃₃₉–₇₁₉. Here we further investigate the validity of AF-predicted conformational ensembles, particularly the preference for the outward-facing W580 conformation in the absence of an allosteric inhibitor. We used MD simulations to explore the effect of NaCl and Trisodium citrate concentrations on the conformational properties of MALT1(PCASP-Ig3)₃₃₉–₇₁₉ under two distinct conditions: first low ionic strength, where electrostatic interactions are pronounced; second high ionic strength, where electrostatic shielding is maximal. Applied in this study an AlphaFold-MD-NMR approach to model the conformational dynamics and activation mechanisms of MALT1provides insights into inter domain movements, and key allosteric regulation events that control MALT1’s catalytic activity.

## 2.0 Result

### 2.1 MD Simulation Study

#### 2.1.1 Dynamics of MALT1(PCASP-Ig3)_339–719_ Under Low-Salt Conditions

The starting structure is critical, as differences in initial conformations can strongly influence the resulting ensemble. To obtain an ensemble of conformations whose back-calculated relaxation parameters best matched the NMR experimental data for MALT1(PCASP-Ig3)_339–719_ measured in buffer containing 20 mM Tris (pH 7.5), 60 mM NaCl, and 1 mM TCEP, we performed free MD simulations starting from a four initial structures (**Table 1**), all based on the sequence used in the NMR study (BMRB accession code 52265).

**Table 1.**
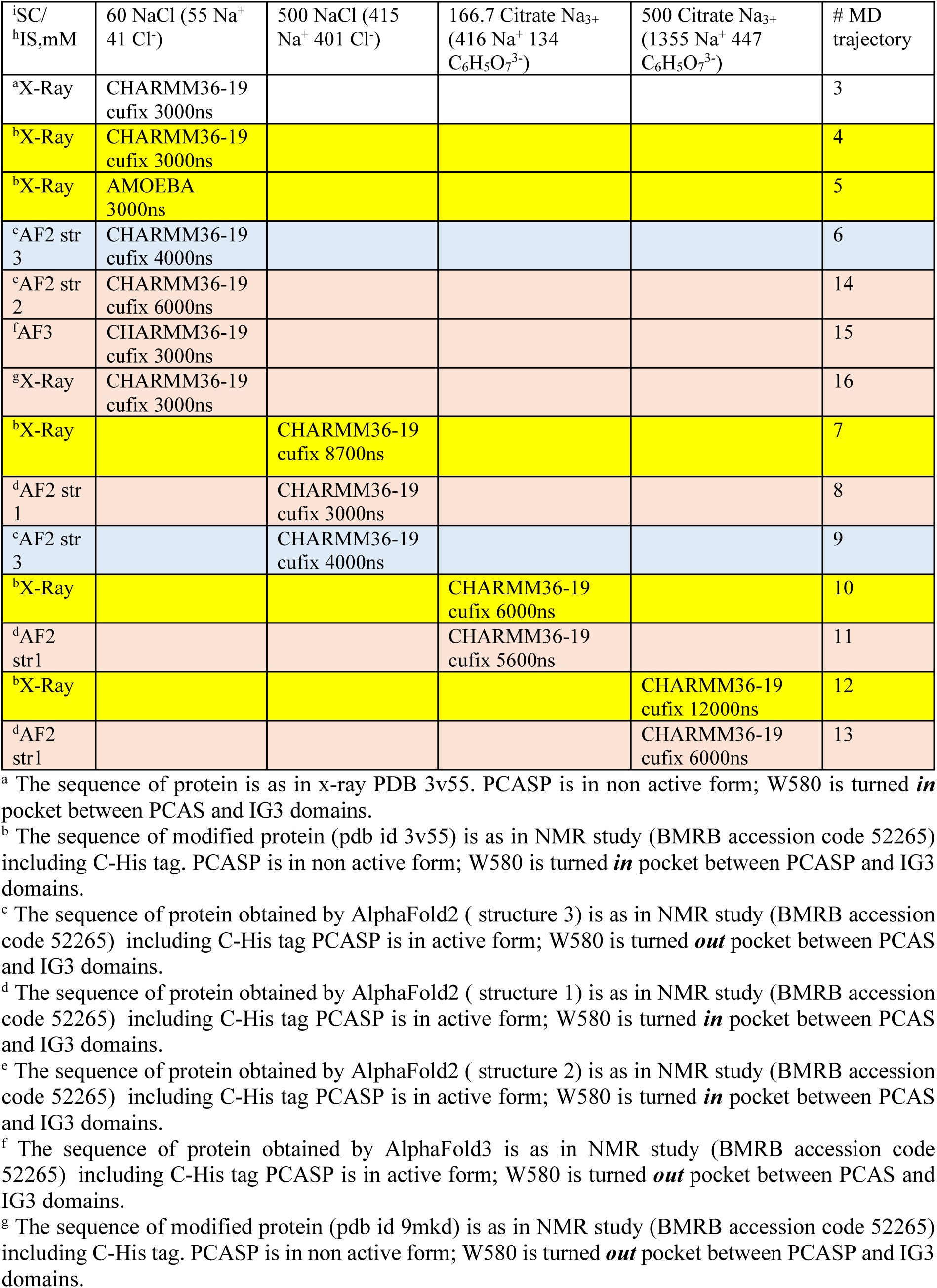
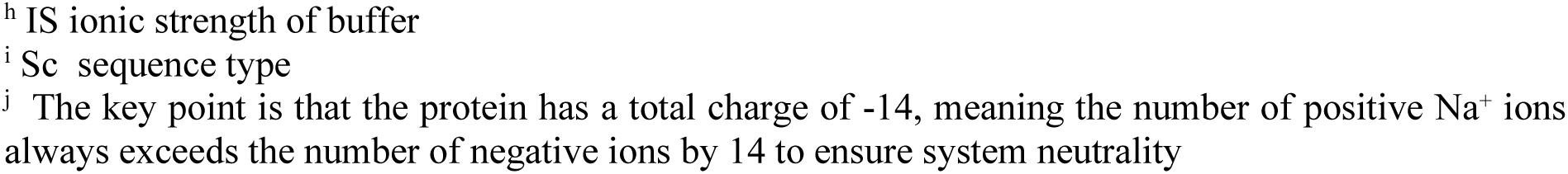
Description of the 1-14 trajectories obtained in free MD simulations^j^.

According to AF, MALT1(PCASP-Ig3)_339–719_ adopts two major structural states distinguished by the orientation of the W580 aromatic ring, which is positioned either within or outside the pocket between the PCASP and Ig3 domains. Throughout the text, these are referred to as **inward-** or **outward-facing W580 conformations,** structures **4, 14** and **6, 15**, respectively. In addition, the PCASP domain may adopt either an inactive or an active conformation, **4, 15** and **6, 14**, respectively. The inactive state resembles the X-ray structure (PDB 3V55), whereas the active state corresponds to the conformation predicted by AF^41^ and resembles those observed in X-ray structures bound to substrate-mimicking ligands (ID: 3V4O).

To assess the quality of the reconstructed trajectories, we evaluated several ensemble properties that describe global structural behaviour, including root-mean-square deviation (RMSD) and Principal Component Analysis (PCA). RMSD is the primary metric used to quantify structural variation within the ensemble. As shown in **Figure 1**, RMSD fluctuations were analysed in four ways using the Cα backbone atoms of MALT1(PCASP-Ig3)_339–719_. RMSD was calculated relative to either the initial structure or the representative structure of the most populated cluster. These analyses were performed for (i) selected residue ranges (5–130, 147–154, 172–225, and 235–380) excluding flexible loop regions, and (ii) all residues except the N– and C-terminal segments (residues 1–4 and 381–388).

**Figure 1.**
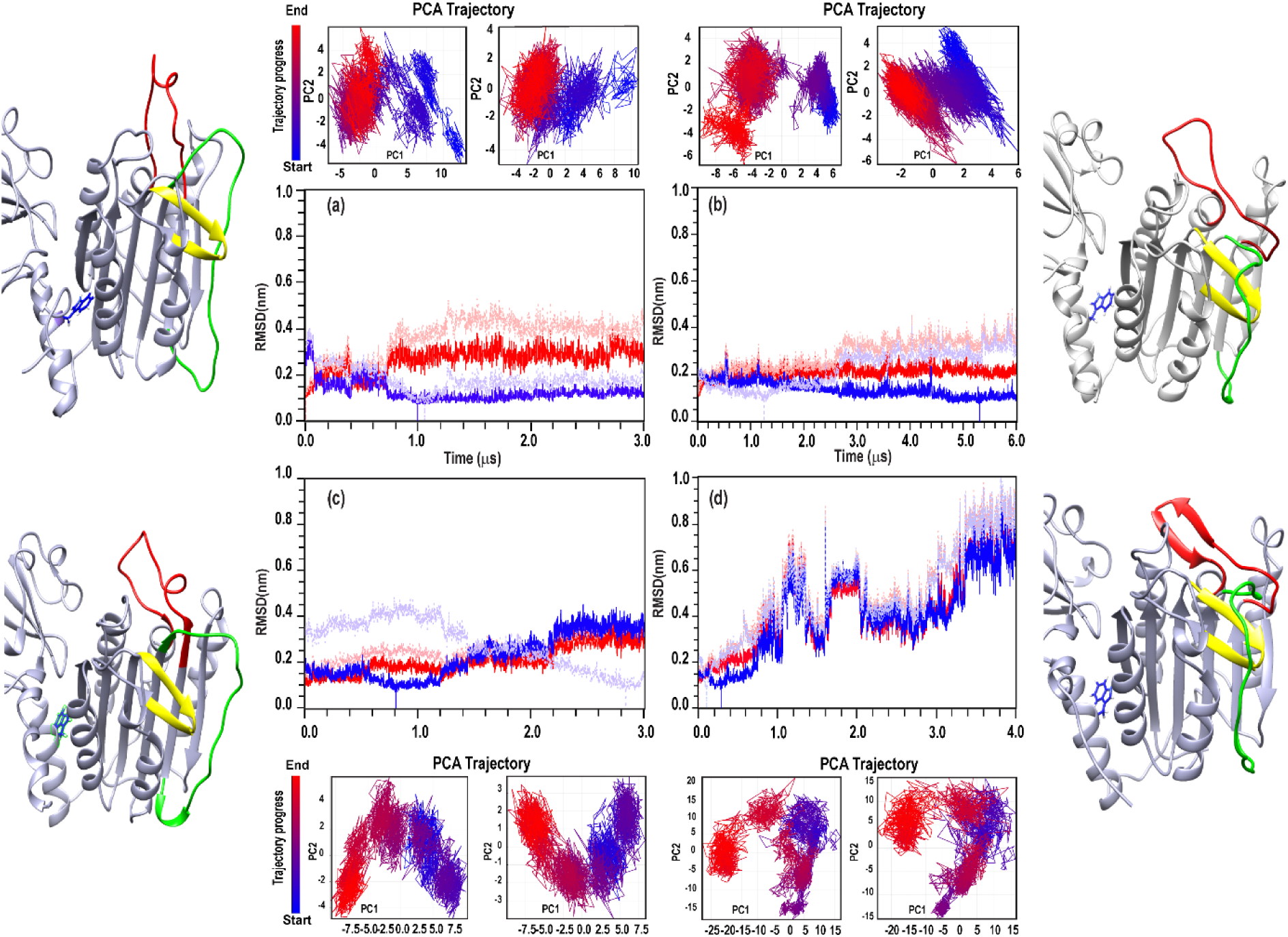
RMSD and PCA of MALT1(PCASP-Ig3)_339–719_ MD trajectories. (a–d) RMSD of Cα atoms relative to the initial structure (red) or the representative structure of the most populated cluster (blue). RMSD was calculated either for selected structured regions 342–467, 484–491, 509–562, and 572–717; dark red/blue) or for all residues except the termini (residues 338–341 and 718–725; light red/blue). Trajectories shown: (a) 4, (b) 14, (c) 15, (d) 6. All simulations were performed under 60 mM NaCl. PCA plots (PC1 vs PC2) are shown above and below each RMSD panel. Left PCA panels use residues 342–717; right panels use structured regions (342–467, 484–491, 509–562, 572–717). Colours from dark highlighting the β3 hairpin (416–425), Loop 2 (464–485), Loop 3 (491–509), and the orientation of W580 (inward or outward, in blue).

PCA is a dimensionality-reduction technique essential for visualizing and characterizing the dominant motions in molecular ensembles, providing a complementary perspective to RMSD for describing global dynamics. To assess whether our models capture these key dynamical features, we analysed the first two principal components (PC1 and PC2) for trajectories **4**, **6**, **14**, and **15**. PCA was performed using two residue selections intervals: residues 5–380 (**Figure 1**, left PCA panels) and residues 5–130, 147–154, 172–225, and 235–380 (**Figure 1**, right PCA panels). The PC1/PC2 projections for individual structures from trajectories **4**, **6**, **14**, and **15** (**Figure 1**) show that the sampled conformations do not simply interpolate between two end states but instead populate a diverse ensemble. However, notable differences are observed among the trajectories.

The most pronounced changes in RMSD and PCA were observed in trajectory **6**, a 4 μs MD simulation initiated from an AF-generated structure (**Table 1**) in which W580 adopts an outward-facing conformation and the PCASP domain begins in an active state. As shown in **Figure 1d**, the RMSD profile of trajectory **6** indicates instability, consistent with an ongoing conformational transition during the MD simulation. Detailed analysis reveals a key structural event: a flip of the W580 aromatic ring involving transitions in the χ₁ and χ₂ dihedral angles (**Figure 2 a, b**). Initially, the χ₁ angle remains stable around −170°, but at ∼800 ns, it shifts to −70°, and by ∼1500 ns, it reverts to −170°. During this interval, the χ₂ angle fluctuates near 100°. However, at ∼2300 ns, a pronounced ring flip occurs, as χ₂ shifts from 80° to −100°. Following this transition, both angles stabilize, and the system adopts an inward-facing W580 conformation (**Figure 2c,d**).

**Figure 2.**
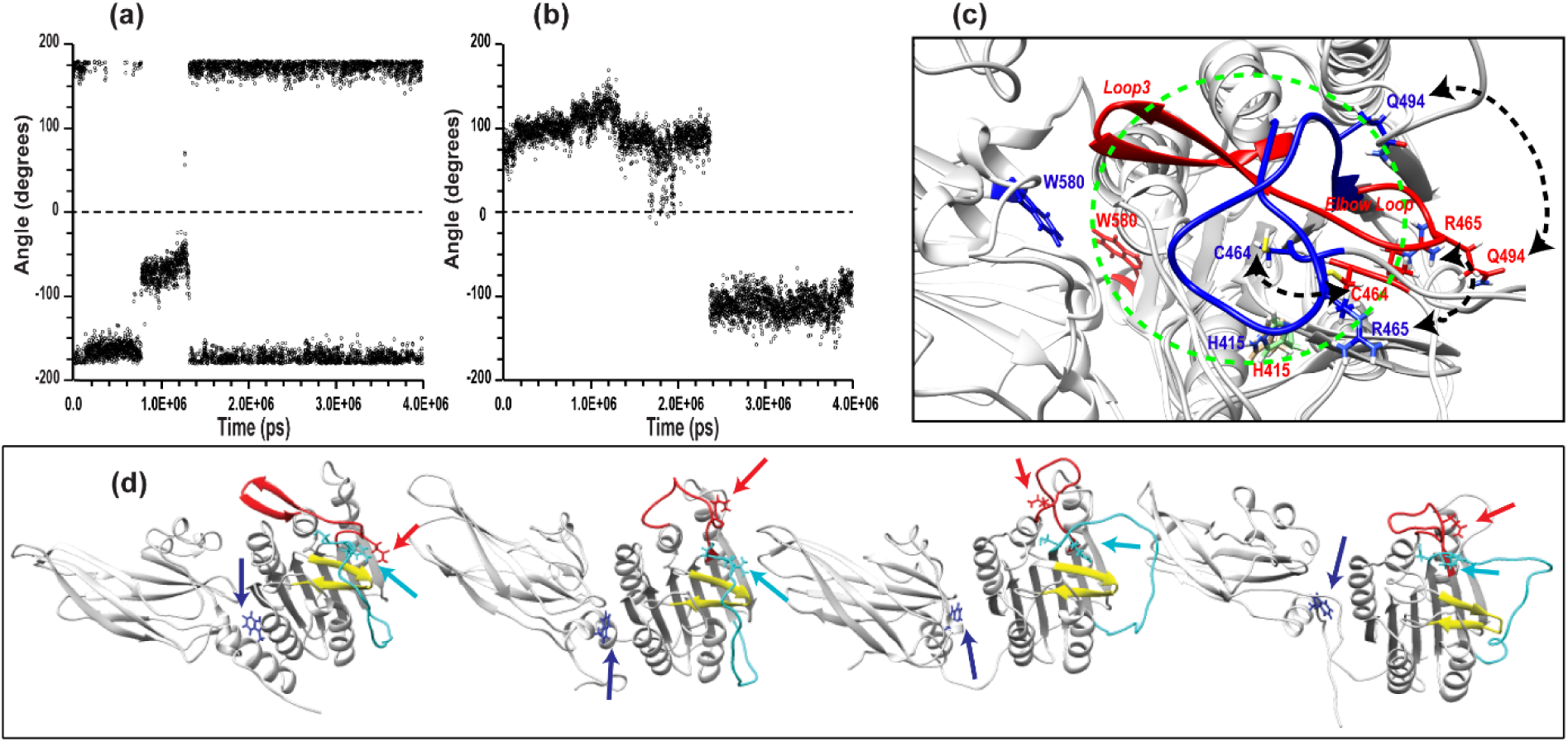
Dynamics of χ₁ and χ₂ Angles Along MD Trajectory 6 of MALT1(PCASP-Ig3)₃₃₉–₇₁₉. (a) χ₁ and (b) χ₂ torsion angles throughout the 4 μs MD simulation, compared to the initial structure of MALT1(PCASP-Ig3)₃₃₉–₇₁₉, which begins in the outward-facing W580 conformation (as described in Table 1). (c) Superimposed structures of MALT1(PCASP-Ig3)₃₃₉–₇₁₉ from the start and end of trajectory **6**, illustrating transitions between the active and inactive states of the PCASP domain. Loop3 and selected amino acids are highlighted in red and blue, respectively, with the rest of the protein shown in grey. The position of the elbow loop (residue 491-498) in the active state is labelled as “Elbow Loop”. Dash lines indicate the displacement of residues C464, R465, and Q494. Two orientations of the **W580** aromatic ring and the position of catalytic H415 are shown. The protein’s active site is marked with a dashed green circle. In the active conformation, this site remains open, while in the inactive state it is occluded by Loop3. (d) Conformational transition of the PCASP domain from active (leftmost structure) to inactive (rightmost structure), highlighting the flip of W580 during a 4 μs MD simulation at low salt concentration. The β3 hairpin loop (residues 416–425), Loop2 (residues 464–485), and Loop3 (residues 491–509) are coloured yellow, cyan, and red, respectively. W580 and Q494 are shown in blue and red, respectively, while R465 and C464 are shown in cyan.

Visual inspection of the PC1/PC2 projections (**Figure 1**) highlights the conformational diversity of trajectories **6**, in which up to four major structural clusters can be distinguished. To further characterize heterogeneity and cooperative motions within the MALT1(PCASP-Ig3)_339–719_ ensembles in trajectory **6** captured by PC1 and PC2, we visualized the corresponding PCA loadings by superposing four representative MD structures that reflect the structural variations along each principal component (**Figure 2c,d**). The transition from an outward-to inward-facing conformation of W580 in MALT1(PCASP-Ig3)₃₃₉–₇₁₉ is shown in **Figure 2d**. This complex aromatic ring flip W580 occurs through a refolding process that involves both the Ig3 and PCASP domains, which move semi-independently, consistent with our previous findings^41^ and as reflected by the continued RMSD changes (**Figure 1d**). A particularly intriguing result from the MD simulations was that the W580 flip seems to be coupled with a rearrangement of the active site in the PCASP domain in Loop3.

The system, which began in the active conformation, transitioned into the inactive state similar to that observed in trajectory **4** and illustrated in **Figure 2c, d**. This transition is driven by coordinated movements of Loop 2 and Loop 3.

A distinct two-site flip of Loop 2 (cyan in **Figure 2d**) across the β3 hairpin loop (residues 416–425, yellow) occurs at approximately 2.5 µs in trajectory **6**, placing Loop 2 in a position similar to that seen in the inactive starting structure of trajectory **4**. This repositioning of Loop 2 subsequently displaces Loop 3 to the top of the MALT1 active site (**Figure 2c**).

This rearrangement is most clearly tracked through a set of conformational “fingerprints” described by Zhang et al.^42^, involving residues C464, R465, E549, and Q494, identified through comparisons of MALT1 crystal structures bound to an allosteric inhibitor (PDB 6F7I) or a peptidic inhibitor (PDB 3V4O).

In our study, trajectory **6** reveals that a distinctive feature of the active conformation like the presence of a short ***elbow Loop***^13, 42^ (residues 491–498) within Loop3 (residues 490–509) (**Figure 2c** shown in red and **2d** leftmost structure) is absent in the inactive state (**Figure 2c** shown in blue and **2d** rightmost structure). In the inactive state, Loop3 folds over the catalytic dyad (C464–H415), effectively blocking substrate access to the active site (**Figure 2c** shown in blue). As a consequence of this rearrangement, we observed significant moves, up to 10 Å for Q494 and up to 6 Å for R465, from active to inactive conformations of PCASP, both of which are closely positioned to E549 in the active conformation.

Similar to trajectory **6**, trajectory **15,** initiated from outward-facing W580 conformations with the PCASP domain the inactive state, shows a dominant tendency for the W580 aromatic ring to flip from an outward-to an inward-facing conformation. This is reflected by unstable (though less pronounced than in trajectory **6** RMSD variation (**Figure 1c**) and by the conformational diversity observed in the PC1/PC2 projections (**Figure 1**). As in of trajectory **6**, up to four major structural clusters can be distinguished in the PC1/PC2 plane for trajectory **15**. The PCA loadings (**Figure 1**) further illustrate the transition of W580 toward an inward-facing conformation in MALT1(PCASP-Ig3)₃₃₉–₇₁₉, although the positions of Loop2 and Loop3 even if flexible remain characteristic of the inactive state.

These results align with our previous free energy calculations^41^, which indicated that the inward-facing W580 conformation is energetically preferred by ∼0.67 ± 0.2 kcal/mol lower in energy than the outward conformation.

In striking contrast to trajectories **6** and **15**, RMSD analysis of MALT1(PCASP-Ig3)_339–719_ trajectory **4** indicates structural stability after 0.8μs MD simulation, with no significant deviations throughout the 3 µs simulation (**Figure 1a**). This behaviour is consistent with the PCA analysis. The conformational evolution in trajectory **4** captured by PC1 and PC2 reveals a small number of low-population states early in the simulation (up to ∼0.8 µs; structures coloured blue) taken place in an equilibration period of MD simulation, after which the system transitions into a single dominant cluster (purple/red) that persists for the remainder of the trajectory. The system, initiated with the inward-facing W580 conformation, remains in the passive form of the PCASP domain of MALT1(PCASP-Ig3)_339–719_ under low-salt conditions (60 mM NaCl).

Finally, we investigated whether a spontaneous transition from the active PCASP conformation to the in-active, inward-facing W580 conformation can be captured in long MD simulations under low-salt conditions (60 mM NaCl). To address this, we performed an additional 6μs MD simulation (trajectory **14**), starting from the AF-generated structure (**Table 1**) in which the PCASP domain was initialized in the active conformation, unlike trajectory **4**, which began in the inactive state, while both shared an inward-facing W580 conformation.

RMSD analysis of the MALT1(PCASP-Ig3)_339–719_ trajectory **14** calculated over backbone atoms excluding flexible loop regions, indicates high structural stability with no major deviations throughout the 6 µs simulation (**Figure 1b**, dark red and blue curves). In contrast, when flexible loop regions are included, a transition to a higher RMSD plateau is observed at approximately 2.5 µs (**Figure 1b**, light pink and grey curves). This behaviour is consistent with the PCA results: the PC1/PC2 projections for trajectory **14** show two dominant clusters (purple/red) that persist for the remainder of the simulation.

Supplementary **Figure S1** shows superpositions of 30 structures sampled from opposite ends of the structural span defined by the PC1 modes for all conformational ensembles. Visual inspection reveals two key features: (i) the inward-facing orientation of the W580 aromatic ring is maintained throughout trajectory **14**, consistent with the behaviour observed in trajectory **4**; and (ii) loop 2 flips around the protruding β-sheet spanning residues 416–426, leading to a transition of the PCASP domain from an active to an inactive conformation.

In summary, all four trajectories, despite starting from different active-site conformations and W580 orientations, converged to a common conformational ensemble characterized by an inward-facing W580 aromatic ring and an inactive PCASP active site in MALT1(PCASP-Ig3)_339–719_ under low-salt conditions (60 mM NaCl).

#### 2.1.2 Dynamics of MALT1(PCASP-Ig3)_339–719_ in 166.7 mM Trisodium Citrate (Kosmotropic Conditions)

As the next step, we used MD simulations to investigate the effect of salt on the conformational properties of MALT1(PCASP-Ig3)_339-719_. Instead of employing physiological electrolytes such as NaCl, we mimicked the *in vitro* assay conditions by using a kosmotropic buffer, trisodium citrate, at concentrations up to 166.7 mM. These conditions correspond to a high–ionic-strength environment in which electrostatic shielding shift the interplay of electrostatic and hydrophobic interactions within the protein.

Two MD calculation were initiated using starting structures as for trajectories **4** and **14** both with as inward W580 conformations, but the PCASP domain may adopt either an inactive or an active conformation, creating trajectories **10** and **11**, respectively.

No significant structural changes were observed in trajectory **10**: both the RMSD profile and the PC1/PC2 projection remained consistent with the conformational ensemble obtained for trajectory **4**. In contrast, trajectory **11** revealed the most intriguing structural dynamics of MALT1(PCASP-Ig3)_339-719_ in 166.7 mM trisodium citrate (**Figure 3**).

**Figure 3.**
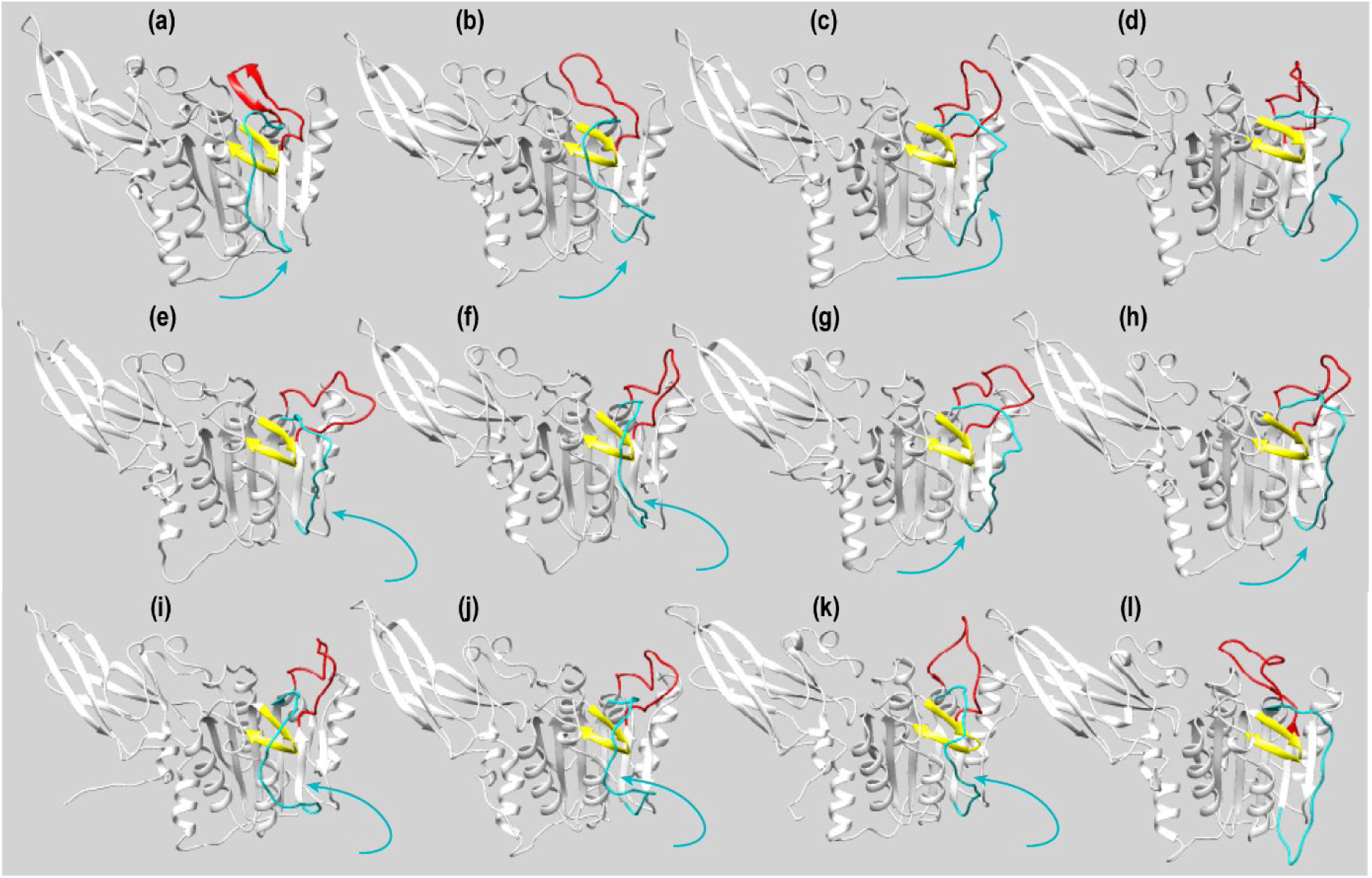
Structural Dynamics Along MD Trajectory 11 of MALT1(PCASP-Ig3)₃₃₉–₇₁₉ at 166.7 mM Trisodium citrate. (a–k) Selected snapshots from MD trajectory 11 showing the conformational transition of Loop2 (residues 464–485, highlighted in cyan) around the β3 hairpin loop (residues 416–425, highlighted in yellow) in MALT1(PCASP-Ig3)₃₃₉–₇₁₉. These frames illustrate the dynamic shift between two distinct conformations of Loop2. The starting structure (a) represents the active conformation of the PCASP domain. Structures (b)–(k) were extracted from the MD trajectory at 300, 1710, 1740, 2220, 2670, 3060, 3120, 3150, 3210, and 3840 ns, respectively. (l) For comparison, a structure of the PCASP domain in the inactive state, modelled based on the crystal structure template PDB ID: 3V55, is shown.

As shown by the MD snapshots, Loop **2** (residues 464–485, highlighted in cyan) undergoes a dynamic equilibrium between two distinct conformations around the β3 hairpin loop (residues 416–425, highlighted in yellow).

Starting from the active-state conformation of the PCASP domain (**Figure 3a**), Loop 2 shifts into the inactive-state position (**Figure 3b–d**), then returns to the active state (**Figure 3e–f**), transitions again to the inactive state (**Figure 3g–h**), and once more to the active state (**Figure 3i–k**), ultimately adopting the inactive conformation at the end of the 3 μs simulation (**Figure 3l**). Notably, this transition is accompanied by conformational fluctuations of Loop **3**, consistent with the behaviour described for trajectory **6** in **Figure 2** during the active-to-inactive transition of the PCASP domain.

The unique dynamic behaviour observed in trajectory **11** suggests that in the kosmotropic trisodium citrate buffer, the equilibrium between inactive and active conformations is slightly shifted toward the active state within the accessible MD timescale. Such a shift was not detected in trajectories **4** (low-salt conditions) or **10** (trisodium citrate buffer), where no inactive-to-active transition occurred even in simulations extending up to 7 μs. These observations imply that the full conformational transition likely occurs on a much slower, millisecond timescale, making it challenging to fully capture within typical MD simulation lengths.

#### 2.1.3 Dynamics of MALT1(PCASP-Ig3)_339–719_ at high salt (500 mM NaCl)

To investigate how ionic strength influences the conformational behaviour of MALT1(PCASP-Ig3)_339-719_, we performed MD simulations under high-salt conditions (0.5 M NaCl), in which electrostatic interactions are maximally shielded. Three simulations, corresponding to trajectories **7**, **8**, and **9**, were initiated from distinct starting structures (**Table 1**).

Trajectories **4** and **7** share the same starting structure, differing only MD simulation in ionic strength. RMSD analysis of trajectory 7 (**Figure S2g**) indicates that the structure remains stable throughout the simulation, with no significant deviation from the initial conformation.

To further probe the behaviour of the PCASP domain under high-salt conditions, we carried out a 3 μs simulation (trajectory **9**), using the same AF-generated model as in trajectory **6**, i.e. with W580 in an outward-facing conformation and an active PCASP domain. RMSD analysis (**Figure 4c**) shows that trajectory **9** remains close to the starting structure. Consistently, W580 retains its outward orientation, and the PCASP active site remains in the active state. The PC1/PC2 analysis likewise shows a single, well-defined conformational family (**Figure 4c1,c2**), in contrast to the broad distribution observed for trajectory **6** (**Figure 4d1,d2**) after the equilibration period (∼0.3 μs).

**Figure 4.**
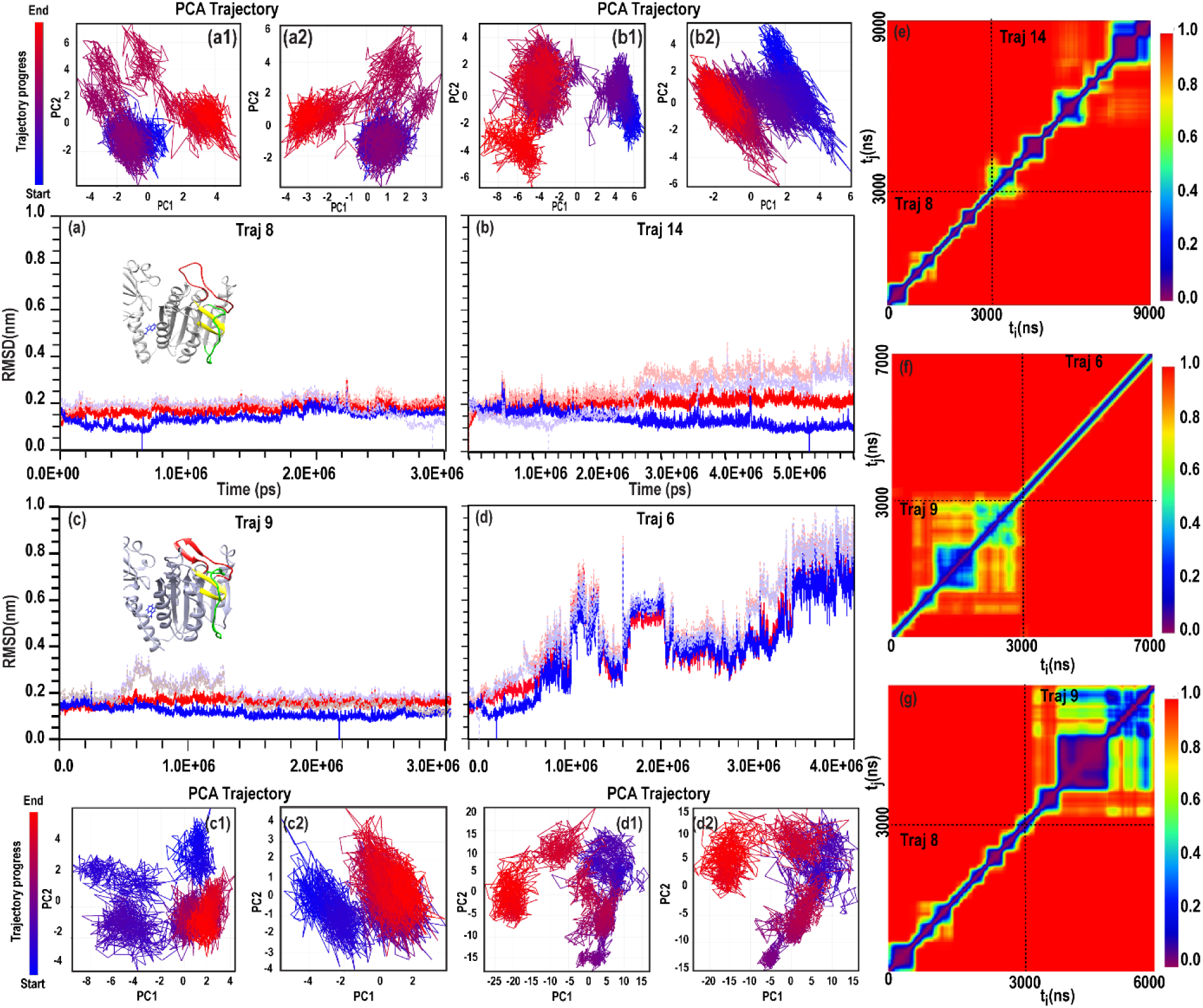
Conformational dynamics of MALT1(PCASP-Ig3)_339-719_ under high ionic strength (0.5 M NaCl). (a–d) Backbone RMSD of trajectories 8 (a), 14 (b), 9 (c) and 6 (d) relative to their respective starting structures (red) and best cluster structure (blue). (a1–a2, b1–b2, c1–c2) Principal component (PC1/PC2) projections of trajectories 8 (a1,a2), 14 (b1,b2), and 9 (c1,c2), showing the distribution of conformational states after equilibration. (d1–d2) PC1/PC2 projections for trajectory 6 following equilibration, illustrating its broad and heterogeneous conformational space. (e) Conformational clustering of the combined trajectories 8 and 14, revealing multiple local minima and non-overlapping conformational families. (f) Conformational clustering of the combined trajectories 9 and 6, showing a single dominant conformational minimum for trajectory 9 and highly dispersed conformations for trajectory 6. (g) Conformational clustering of the combined trajectories 9 and 8, demonstrating that trajectory 9 populates a single stable local minimum, whereas trajectory 8 samples several small, unstable clusters.

A striking difference emerges when the 7 μs combined trajectory (trajectories **9** + **6**) is subjected to conformational clustering (see Methods). As shown in **Figure 4f**, trajectory **9** occupies a single, well-populated local minimum, whereas trajectory **6** displays no stable clusters, consistent with its higher conformational heterogeneity. The two trajectories populate non-overlapping regions of conformational space.

We next performed a 3 μs simulation (trajectory **8**) starting from the same AF-generated structure used for trajectory **14**, in which W580 is inward-facing and the PCASP domain adopts an active conformation. Unlike trajectories **6** and **9**, which show pronounced differences in structural drift, the RMSD profiles of trajectories **8** and **14** are more similar and less conclusive. It seems that RMSD analysis of trajectory **8** (**Figure 4a**) indicates a stable structure with no major deviations from the starting conformation. Structural inspection confirms that W580 remains inward-facing and that the active site of the PCASP domain is preserved throughout the simulation.

PC1/PC2 analysis shows two well-defined conformational families in trajectories **8** and this in also found in trajectory **14** (**Figure 4a1, a2** and **4b1, b2**). However, trajectory **14** undergoes a transition from the active to the inactive PCASP conformation, whereas this transition is not observed in trajectory **8**. Clustering of the combined trajectories **8** + **14** (**Figure 4e**) reveals several local minima, again indicating conformational heterogeneity. The clusters populated by the two trajectories are non-overlapping.

An unexpected and notable result is that trajectory **9** is more structurally stable than trajectory **8** under high-salt conditions, as demonstrated by clustering of the combined trajectories (**Figure 4g**). Trajectory **9** occupies a single, well-populated local minimum, while trajectory **8** populates several small, unstable clusters. No overlap is observed between their respective cluster distributions.

In summary, under high ionic strength (0.5 M NaCl), W580 samples both inward– and outward-facing orientations, with an apparent preference for the latter, unlike in low-salt conditions, where only the inward conformation is observed. Moreover, none of the high-salt simulations show a transition to the inactive PCASP state.

#### 2.2.1 Comparison of MD ensembles with experimental backbone relaxation in MALT1(PCASP-Ig3)_339–719_

To evaluate the conformational ensembles generated by MD simulations, we back-calculated backbone ¹⁵N relaxation parameters (R₁, R₂, and ¹H–¹⁵N NOE) and CSA/dipole cross-correlation (η_xy_) rate from trajectories **4**, **8**, and **9** (**Table 1**) and compared them to experimental NMR data obtained under low-salt conditions. We analysed both R₂ and η_xy_ rates because R₂ can be significantly influenced by slow exchange dynamics, which current back-calculation methods do not account for, potentially introducing inaccuracies. In contrast, η_xy_ is insensitive to slow exchange and can be accurately predicted from MD^43^. Trajectory 4 maintained the PCASP domain in its inactive state with W580 positioned in the interdomain pocket, while trajectories **8** and **9** retained the active PCASP conformation with W580 inward– or outward-facing, respectively. All trajectories were stable, and the 2500–3000 ns segment from trajectory was used for back-calculation.

Figure 5 shows a zoomed region of MALT1(PCASP-Ig3)_339–719_ highlighting Loop 2, Loop 3, and Loop 5. R₁ and η_xy_ values were back-calculated from 500 ns segments of trajectories **3**, **4**, and **8** and compared to experimental data acquired at 900 MHz. The main differences between trajectories **4** and **8**, **9** and experimental data occur in disordered segments spanning residues 496–510 (Loop 3) and 566–580 (Loop 6). (**Figure S3**).

**Figure 5.**
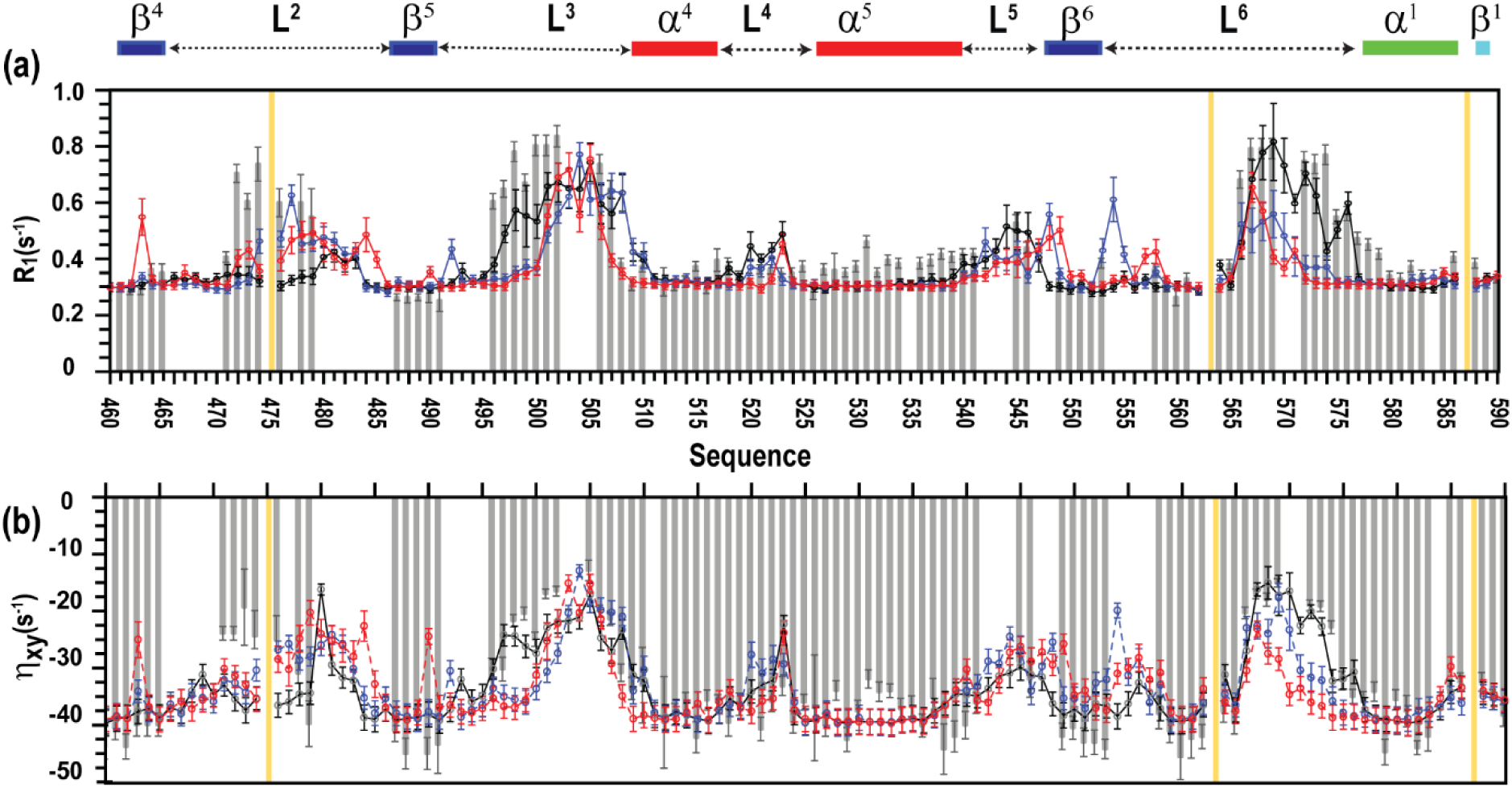
Dynamic parameters of the MALT1(PCASP-Ig3)_339–719_, amide backbone obtained on 900 MHz spectrometers. The relaxation parameters for residues 460-590 of the MALT1(PCASP-Ig3)_339–719_ are shown: (**a**) longitudinal relaxation rate R_1_ (s^−1^), and (**b**) CSA/dipole cross-correlation relaxation rates, η_xy_. Experimentally measured R_1_ and η_xy_ values are shown as light grey solid brackets. The corresponding parameters predicted from molecular-dynamics ensembles **8**, **9** and **4** are shown as solid blue, red and black lines, respectively. Ensembles 8 and 9 were generated at high salt concentration from trajectory segments 2500–3000 ns and 3500–4000 ns, respectively, using the CHARMM36-19 force field. Ensemble 4 was generated at low salt concentration from trajectory segments 2500–3000 ns using the same force field. Experimental error bars represent one standard deviation from curve fitting; predicted parameter uncertainties were estimated by bootstrap analysis.

To quantify agreement between experimental and calculated R₁, R₂, ¹H–¹⁵N NOE and η_xy_ values, two test were applied. First, a cosine-based criterion (equation 7 in Method) was used to rank trajectories according to the lowest score. Additionally, the RMSE test was applied to the best-scoring trajectory. Complete results from the cosine ranking and RMSE tests for all trajectories are presented in **Table 3**. Overall, trajectory **4** obtained the lowest score and reproduces experimental R₁, R₂, ¹H–¹⁵N NOE and η_xy_ most closely, compare with trajectories 8/9.

In summary, these results indicate that MD ensembles initiated from the NMR-based structure (trajectory **4**) provide the most accurate representation of backbone dynamics for MALT1(PCASP-Ig3)_339-719_ and capture the global conformational space under low-salt buffer conditions.

#### 2.2.2 Comparison of MD ensembles created from AF and X-ray structures of in MALT1(PCASP-Ig3)_339–719_

Next, we examined whether the conformational ensemble sampled in trajectory **4**, initiated from the AF-derived sequence, is consistent with ensembles generated from X-ray–based starting models of inactive MALT1. To this end, we analysed three MD trajectories of MALT1(PCASP-Ig3)_339–719_, under identical buffer conditions (20 mM Tris pH 7.5, 50 mM NaCl, 1 mM TCEP), differing only in their initial structures.

Trajectory **3** was initiated from the X-ray structure (PDB 3V55), which includes five additional N-terminal residues, whereas trajectory **4** used the sequence from our NMR study (BMRB 52265) with a six-residue C-terminal His-tag.

A third trajectory **16** employed the recently solved inactive X-ray structure (PDB 9MKD)^44^, where Loop 3 adopts a partially α-helical conformation and helix α5 is entirely disordered.

All starting structures placed the PCASP domain in the inactive state. In trajectories **3** and **4**, W580 occupies the interdomain pocket, while in trajectory **16** it adopts an outward-facing orientation. Terminal extensions were treated as disordered regions.

In addition to described above relaxation rates, the relaxation back calculated for trajectory **4**, the corresponding values were calculated for trajectories **3** and **16**. Backbone relaxation parameters (R₁, R₂, η_xy_, and NOE) were back-calculated from 500 ns segments (2500–3000 ns) of all trajectories, **3**, **4**, **16** and compared with experimental data (Figures 6 and **S4**).

**Figure 6.**
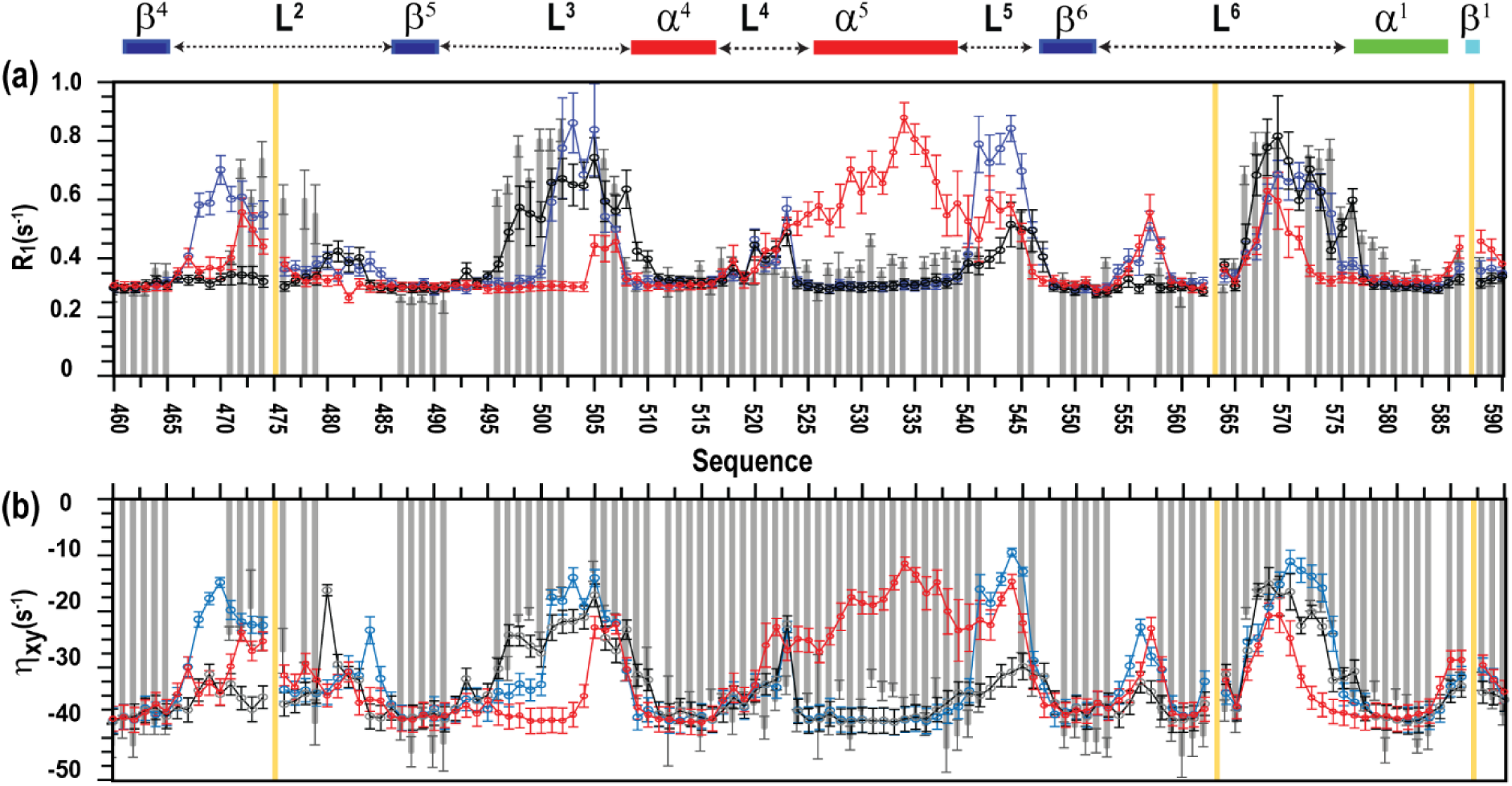
Dynamic parameters of the MALT1(PCASP-Ig3)_339–719_, amide backbone obtained on 900 MHz spectrometers. The relaxation parameters for residues 460-590 of the MALT1(PCASP-Ig3)_339–719_ are shown: (**a**) longitudinal relaxation rate R_1_ (s^−1^), and (**b**) CSA/dipole cross-correlation relaxation rates, η_xy_. Experimentally measured R_1_ and η_xy_ values are shown as light grey solid brackets. The corresponding parameters predicted from molecular-dynamics ensembles **3**, **4** and **16** are shown as solid blue, red and black lines, respectively. Ensembles 3, 4 and 16 were generated from trajectory segments 2500–3000 ns using the CHARMM36-19 force field at low salt concentration. Experimental error bars represent one standard deviation from curve fitting; predicted parameter uncertainties were estimated by bootstrap analysis.

Initial inspection shows that trajectories **3** and **4** differ only modestly, with minor deviations confined to a few residues in Loop 2, which appears slightly more flexible in trajectory **3**. In contrast, ensemble **16** exhibits pronounced discrepancies relative to experiment, most notably across the α6 region, where the predicted dynamics diverge substantially from the measured values. These observations are quantitatively supported by the cosine-similarity and RMSE analyses (**Table 3**), which rank trajectories **3** and **4** as most similar to experiment, whereas ensemble **16** yields the poorest scores, reflecting its significant deviation from the experimentally observed dynamics.

In summary, these results show that the MD ensemble initiated from the AF/NMR-derived structure (trajectory **4**) provides the most accurate representation of MALT1(PCASP-Ig3)_339-719_ backbone dynamics and closely agrees with the ensemble generated from the inactive X-ray structure (PDB 3V55).

### 2.3 Methyl Group Dynamics as Probes of Conformational Stability: Agreement Between MD and NMR

Overall, the methyl longitudinal relaxation rate (R₁) and cross-correlated relaxation rate (Г₂), (**Figure S5**) provide a consistent benchmark for evaluating MD-derived dynamics. In our previous study on the apo form of human MALT1(PCASP-Ig3)_338–719_^40^ we reported nearly complete^1^H /^13^C methyl resonance assignments for Ile, Leu and Val residues. Trajectories ensembles, **4** (low salt) and **8**, **9** (high salt), closely reproduce the experimental relaxation parameters across Ile, Leu, and Val residues (**Figure S5a–f**), indicating that the MD simulations accurately capture fast side-chain motions. More thorough analyses of the methyl dynamic of the MALT1(PCASP-Ig3)_338–719_ as well as exponential data for R₁ and (Г₂) are presented in Supplementary.

Methyl relaxation analysis (R₁ and Г₂; **Figure S6**) reveals that the assigned Ile, Leu and Val methyl groups segregate into four well-defined hydrophobic clusters (Cl1–Cl4). Cl1 is located within the IgL3 domain, Cl2 resides at the Ig3–PCASP interface, and Cl3–Cl4 occupy opposite sides of the PCASP β-sheet (**Figure. S6f**).

For Cl1, which lies far from the structural differences between ensembles **4** and **8**, **9** back-calculated R₁ and Г₂ values agree closely with experiment (**Figure S6 d1, d2**). Clusters Cl3 and Cl4 similarly show only minor variation between ensembles (**Figure S6 a1, a2; b1, b2**), indicating that Loop 3 rearrangements within the PCASP domain exert little influence on fast side-chain dynamics in the β-sheet. Surface-exposed methyl’s (**Figure S6 e1, e2**) also display excellent agreement with experiment under both salt conditions.

Cl2, positioned at the Ig3–PCASP interface and previously implicated in allosteric ligand binding, remains the most functionally significant cluster^41^. Nevertheless, the relaxation parameters for Cl2 methyl’s (**Figure S6 c1, c2**) are nearly identical across ensembles **4**, **8** and **9**, despite their distinct active-site conformations.

Together, these observations indicate that the hydrophobic clusters act as stabilizing elements that preserve local fast dynamics even as the protein undergoes larger-scale conformational transitions. The minimal impact of the active ↔ inactive switch on methyl dynamics underscores the rigidity of these clusters and their role in maintaining the overall architecture of MALT1(PCASP–Ig3).

### 2.4 Backbone Dynamics of MALT1(PCASP–Ig3)₃₃₉–₇₁₉ Ensembles

To investigate residue-level dynamics in the MALT1(PCASP–Ig3)₃₃₉–₇₁₉ ensembles, we applied two complementary approaches: analysis of Cα root-mean-square fluctuations (RMSF) and evaluation of the NH-vector autocorrelation function, acf(t). First, we analysed per-residue RMSF profiles to assess regions of structural flexibility across the MD trajectories of MALT1(PCASP–Ig3)₃₃₉–₇₁₉.The averaged RMSF values for trajectories **4**, **8** and **9** are shown in Figure 7a. Overall, residues 560–720 remained relatively stable in all simulations, with fluctuations below ∼2.5 Å. The major differences between trajectories were localized to Loop 2, Loop 3, and the linker region connecting the PCASP and Ig3 domains (Loop 5).

**Figure 7.**
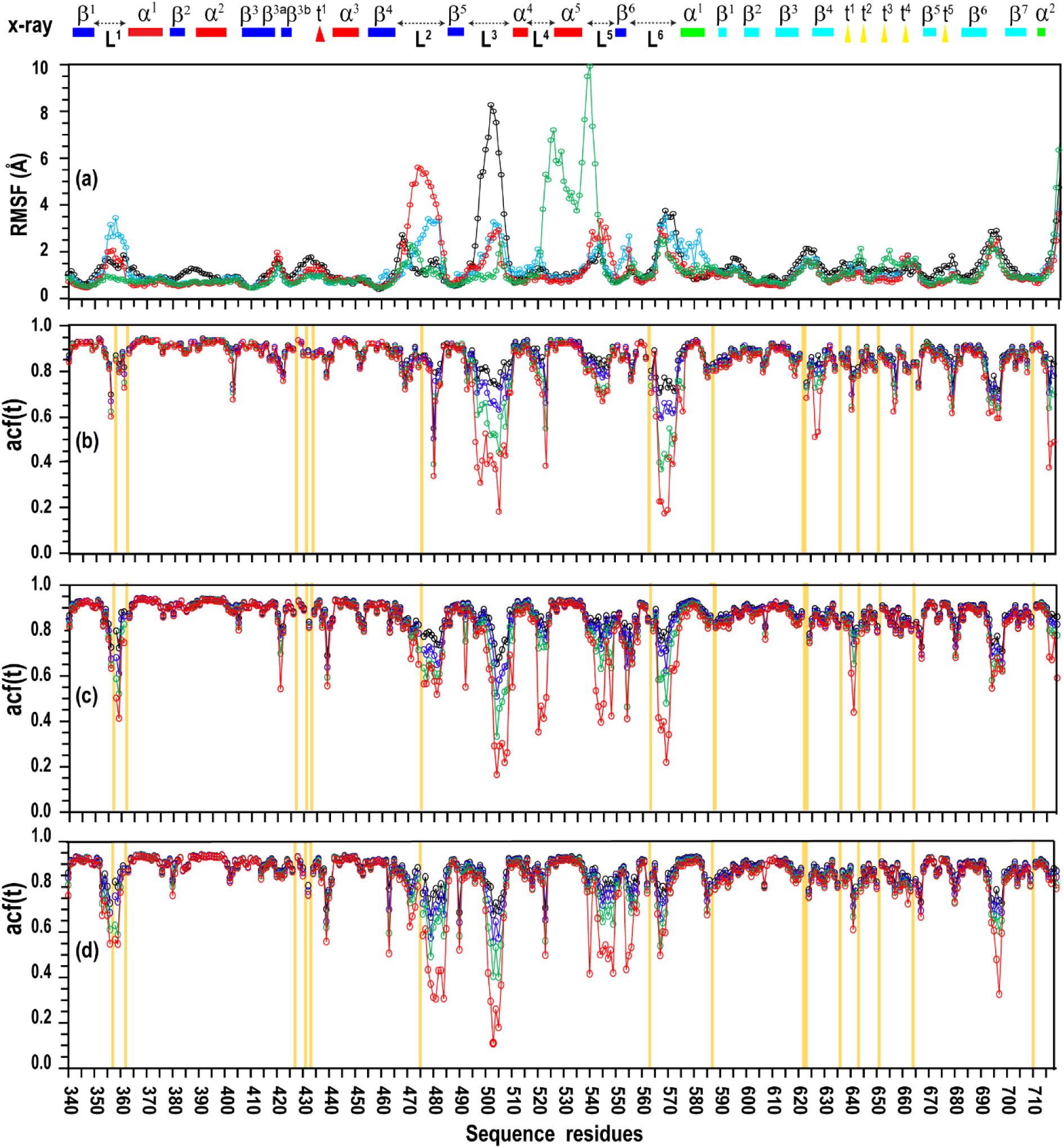
Backbone Dynamics of MALT1(PCASP–Ig3)₃₃₉–₇₁₉ Ensembles. (a) The average RMSF of ensemble of trajectory 4 (black), trajectory 8 (blue), trajectory 9 (red) and trajectory 16h (green) are represented in a line chart. Secondary structure is presented above panel. (b)-(d) Autocorrelation function values, acf(t), were calculated for each residue of MALT1(PCASP-Ig3)₃₃₉–₇₁₉ from the NH vector autocorrelation function at *t* = 0.1, 1.0, 10, and 100 ns (black, blue, green, and red curves, respectively). Data were extracted from the corresponding segment of the MD simulations are shown for trajectories **4** (b), **8**(c) and **9**(d). Secondary structure elements from the crystal structure (PDB 3V55) are indicated above the plots: in the PCASP domain, β-strands (dark blue boxes), α-helices (red boxes), and loops (dashed arrows); in the Ig3 domain, β-strands (light blue boxes), α-helices (green boxes), and short turns (yellow triangles). Proline residues in the protein sequence are indicated by yellow boxes in panels (b) – (d).

In trajectory **4**, which represents the inactive ensemble, Loop 3 exhibits the largest motions, with RMSF values exceeding 8 Å, whereas Loop 2 remains highly ordered (RMSF < 2.5 Å). The pronounced flexibility of Loop 3 may reflect a prerequisite dynamic state that enables its eventual transition toward the active conformation.

In trajectory **9**, corresponding to a stabilized active conformation with W580 rotated outward, Loop 2 shows the greatest flexibility (RMSF up to ∼6 Å), consistent with allosteric pressure toward the inactive Loop 2 position despite the overall rigidity imposed by high-salt conditions. By contrast, trajectory **8**, active-like but with W580 already oriented inward, exhibits intermediate fluctuations across all loops, suggesting that this ensemble samples multiple local minima within the active-state basin.

Interestingly, Loop 1, located near the active site, remains relatively stable in all simulations (RMSF < 2.5 Å), with only a slight increase in flexibility observed in Loop 8 (RMSF ≈ 3 Å).

In contrast to RMSF analysis, which reports the average spatial displacement of Cα atoms across the simulated conformational ensemble, the NH-vector autocorrelation function, acf(t), provides time-resolved information on backbone angular stability. Specifically, acf(t) describes how long the orientation of each NH bond vector remains correlated, thereby revealing the persistence of local structural order over different timescales.

To assess these backbone-motion timescales, we calculated the NH-vector autocorrelation values for every residue of MALT1(PCASP–Ig3)₃₃₉–₇₁₉ at four representative time points: t = 0.1, 1.0, 10, and 100 ns. The resulting acf(t) profiles, extracted from the corresponding MD segments, are shown in Figure 7 for trajectory **4** (panel b), trajectory **8** (panel c), and trajectory **9** (panel d).

As expected, all trajectories show minimal time-dependent changes in acf(t) for residues 340–460, 570–690, and 700–720, with values remaining in the range of 1.0–0.8. This indicates that the NH vectors in these regions are highly stable and dominated by fast picosecond-scale motions. In contrast, the largest deviations and the strongest time-dependent decay in acf(t) are observed in Loops 1–6, with Loops 2, 3, and 6 reaching values as low as ∼0.2 at the 100 ns time point. This pronounced decay suggests the presence of slower motional components within the 500 ns window from which the autocorrelation functions were calculated. Because acf(t) does not reach a plateau for these loop regions, it is likely that additional slow dynamical processes occur on timescales longer than 500 ns.

Together, the RMSF and autocorrelation analyses show that the structured core of MALT1(PCASP–Ig3)₃₃₉–₇₁₉ remains highly stable across all ensembles, whereas backbone dynamics are concentrated in the loop regions. Loop 2 and Loop 3 exhibit the strongest conformation-dependent motions, with Loop 3 most flexible in the inactive ensemble and Loop 2 most flexible in the active ensemble, reflecting their distinct roles in the inactive-to-active transition. Autocorrelation profiles further reveal that these loops undergo slow, long-timescale motions that extend beyond the 500 ns analysis window, whereas the remaining structured regions are dominated by fast picosecond dynamics. Overall, the data indicate that the loop dynamics, rather than the domain cores, define the conformational plasticity of MALT1 and govern its transitions between inactive and active states.

## 3.0 Discussion

MALT1 is a large, multidomain protein whose diverse biological activities depend on the coordinated motions of all its five domains. Even the truncated construct studied here, MALT1(PCASP-Ig3)_339–719_, retains a complex architecture: the Ig3 and PCASP domains form an interconnected framework of α-helices, β-strands, linkers, and loops. Although the Ig3 – PCASP arrangement appears static in the available crystal structures^13^ our earlier MD simulations showed semi-independent motions pivoting around W580, with α-helix connected these two domains acting as a primary driver^41^.

In this study, we focused on loop dynamics in MALT1(PCASP-Ig3)_339–719_, and how they contribute to domain-level transitions.

NMR relaxation data of MALT1(PCASP-Ig3)_339–719_ (R₁, R₂, NOE and η_ex_ profiles) show loop-specific motional difference with much higher flexibility of loop 3 compared to loop 1 (**Figure S3, S4, S9 and S10**), motivating a detailed analysis of loop dynamics and their relationship with domain transition. To capture these motions across timescales, we integrate X-ray structures, AlphaFold (AF) models, NMR relaxation measurements, and long MD simulations.

AF was developed to predict static resolution protein structures^45, 46^, but its ability to sample conformational ensembles or reproduce environmental effects remains limited^41, 47–52^. This limitation is particularly relevant for MALT1, where AF2 and AF3 predict a set of PCASP domain conformations in MALT1(PCASP–Ig3)₃₃₉–₇₁₉ without revealing how structural–functional correlations depend on the surrounding environment.

Both protein conformational equilibria, enzyme activity and ligand binding are sensitive to ionic strength^53–55^, and MALT1 activity itself varies with salt concentration^56^. Although cytosolic ionic strength corresponds to roughly 120–150 mM NaCl equivalent^57^, our initial NMR measurements were performed at 60 mM NaCl. Such sub-physiological ionic strength remains biochemically relevant because it preserves protein stability while allowing detection of conformational states that may be suppressed at higher salt^58, 59^. Given the sensitivity of MALT1 to ionic strength and the sub-physiological salt used in initial NMR experiments, we examined loop behavior under multiple salt conditions.

To minimise structural bias simulations were initiated from both AF-derived starting models and experimentally determined X-ray structures. A set of AF2 and AF3 models, which capture the characteristic inward/outward rotation of W580^41^ and active/inactive-like conformations were used. These were complemented with two crystallographic references: the inactive MALT1 conformation with W580 buried in the interdomain pocket (PDB 3V55)^13^ and an alternative inactive state in which W580 is displaced outside the pocket (PDB 9MKD)^44^.

### Low salt MD calculation of MALT1(PCASP-Ig3)_339–719_

At low-salt conditions (60 mM NaCl), all four AF-initiated trajectories converged to the same inactive inward-facing W580 ensemble, regardless of their initial active-site geometry or W580 orientation. This consistent convergence indicates that reduced ionic strength favours the inactive ensemble for this construct.

A central structural feature of this transition is the coordinated rearrangement of Loop 2 and Loop 3. Their coupled motion aligns with broader principles of loop interdependence in enzyme regulation, where the displacement of one loop can propagate structural change to another^60^. Although Loop 2 (residues 463–485) is often poorly resolved in inactive apo X-ray structures, consistent with intrinsic flexibility, its conformation is well defined in active-state inhibitor-bound structures (e.g., PDB 3UO8, 3V4O, 3V4L, 3UOA, 4I1P, 7PAV, 7PAW).

Prior work by Zhang et al.^42^, comparing PDB 3V4O and 6F7I, already demonstrated that Loop 2 undergoes a substantial positional shift, with residue L480 moving from ∼14 Å (active) to ∼3 Å (inactive) relative to L446 on the α-helix, reinforcing the idea of a major structural transition associated with activation state.

Our simulations capture this coordinated transition of Loop 2, Loop 3 and to the inward rotation of W580, which adopts a stable resting position between the Ig3 and PCASP domains. The trajectories differ only in kinetics: in trajectory **6**, these coupled transitions occur rapidly after equilibration and are accompanied by broader refolding of the protein core, whereas in trajectory **14**, where W580 is already inward facing, the loop transitions proceed more gradually (Figure 1).

Taken together, the MD results indicate that multiple structural blocks surrounding Loop 2, the catalytic-site region, Loop 3, the central PCASP fold, and the PCASP–Ig3 interface, shift coherently as the protein relaxes into its inactive form. The inward positioning of W580 appears to stabilise and promote this collective transition, highlighting its role as a key structural determinant guiding the protein toward an inactive, low-salt–favoured conformation.

In active, inhibitor-bound X-ray structures of MALT1 (e.g., PDB 3V4O and 3V4L), Loop 2 is well ordered, positioned above the β3a–β3b β-hairpin and packed against the α3 helix, thereby stabilizing a correctly aligned catalytic dyad. By contrast, in inactive, ligand-free, or allosterically inhibited conformations (e.g., PDB 3V55, 6F7I), Loop 2 is often unresolved, reflecting disorder. Our MD simulations recapitulate opposite behaviour. In trajectory **14**, the initially active-like Loop 2 samples multiple substates, highlighting intrinsic plasticity that likely primes it for transition.

Once the switch to the inactive state occurs, the MD structure (e.g., trajectory **4**) shows Loop 2 repositioned on the opposite side of the β3a–β3b β-hairpin, close to the β6 strand and to the misaligned catalytic residues. In this conformation, Loop 2 blocks the path normally taken by Loop 3 during activation, preventing Loop 3 from extending toward the β6 strand and keeping the catalytic cleft inaccessible. RMSF and ACF analysis confirm the stability of Loop 2 in this inactive arrangement.

Loop 3 exhibits great structural variability in X-ray structures of inactive MALT1, where it can form a short β-strand (PDB 3V55), an α-helix (PDB 9MKD), or be entirely unresolved. Our MD trajectories similarly show that Loop 3 undergoes substantial fluctuations, yet consistently covers the catalytic cleft in inactive conformations, preventing substrate access.

Importantly, the simulations reveal a defined sequence of activation-related motions: (1) Loop 2 must first move away from its inactive position near the β3a–β3b β-hairpin. (2) This displacement creates the space required for Loop 3 to retract toward the β6 strand. (3) The retraction of Loop 3 opens the active site for substrate binding.

These observations indicate that Loop 3 alone cannot adopt a productive, open conformation without the preceding conformational rearrangement of Loop 2, highlighting a coordinated mechanism underlying MALT1 activation.

### MD calculation of MALT1(PCASP-Ig3)_339–719_ at 166mM salt condition

The consistent collapse of all low-salt trajectories into a single inactive-like ensemble, without any detectable spontaneous return to the active state, suggests that the active conformation of MALT1(PCASP–Ig3)₃₃₉–₇₁₉ is sparsely populated and separated from the inactive state by a substantial kinetic barrier, likely on the millisecond timescale.

This raises the broader question of whether ionic strength modulates the free-energy landscape governing these transitions. When simulations were performed under more physiological electrolyte conditions at salt concentrations up to 166.7 mM, a qualitatively different dynamical behaviour emerged: trajectory **11** displayed a reversible exchange of Loop 2 between its active-and inactive-state positions around the β3 hairpin, accompanied by corresponding fluctuations in Loop 3.

Such behaviour, absent in both the low-salt simulations and those conducted in trisodium citrate buffer, implies that salt composition and ionic strength can subtly reshape the conformational equilibrium of the PCASP domain. In particular, the dynamics observed under the kosmotropic trisodium citrate conditions suggest a modest shift toward the active ensemble within the MD-accessible timescale, hinting that solution environment may play a more active role in tuning MALT1’s catalytic readiness than previously appreciated.

### MD calculation of MALT1(PCASP-Ig3)_339–719_ at high salt condition

Because *in vitro* MALT1 is measurable only at non-physiological, highly kosmotropic conditions^14^, it was essential to examine how elevated ionic strength influences the conformational landscape of MALT1(PCASP-Ig3)_339-719_.

MD simulations at high-salt concentration (0.5 M NaCl or trisodium citrate) revealed a striking stabilization of the starting conformations: W580 remained locked in its initial orientation, Loop 2 and Loop 3 showed markedly reduced fluctuations, and no transitions between active and inactive PCASP states were observed.

Although the overall behaviour was similar for both salts, contrary to that reported in biochemical assays, this discrepancy underscores the need for improved salt parametrization in MD force fields. Nonetheless, the simulations clearly indicate that high ionic strength dramatically suppresses the loop dynamics that facilitate active–inactive transitions, implying that elevated salt concentrations strongly enrich the active-state population of the PCASP domain.

### MD-NMR comparison

Across all MD trajectories of monomeric, ligand-free MALT1(PCASP–Ig3)₃₃₉–₇₁₉, coordinated rearrangements of Loop 2, Loop 3, and the inward rotation of W580 were consistently observed, indicating that these transitions can occur independently of substrate or inhibitor binding. However, because MD samples accessible conformational space rather than experimentally populated ground states, the precise location and stability of the lowest-energy conformers should ultimately be validated by experiment.

Trajectory length and segment selection are critical for back-calculating NMR relaxation parameters, as incomplete sampling can lead to discrepancies with experimental data. This issues have been widely discussed among researchers, including our group^43, 61^ and others^62^ as well in this study. For reliable predictions of ¹⁵N and ¹³C relaxation from the back-calculated from an MD trajectory, the autocorrelation function should be calculated over at least seven rotational correlation times, τ_c_.

To test the adequacy of classical force fields, we compared CHARMM36m and polarizable AMOEBA simulations starting from the same structure used to create trajectory 4 identified as the best agreement with NMR relaxation data. Under low-salt conditions, AMOEBA did not improve agreement with NMR relaxation data, indicating that CHARMM36m sufficiently captures the relevant loop and domain dynamics for this system.

We further incorporated CSA/dipole cross-correlation relaxation (η_xy_), which is insensitive to slow exchange than R₂, and therefore provides a more accurate assessment of loop dynamics, including potential contributions in L3 and L6 that cannot be excluded.

Comparison of experimentally measured relaxation parameters of back bone (R₁, R₂, η_xy_, and NOE) at low-salt shows that MD ensembles initiated from the NMR-derived structure (trajectory **4**) provide the closest agreement with experiment. This trajectory offers the most faithful representation of MALT1(PCASP-Ig3)₃₃₉–₇₁₉ backbone dynamics and most accurately captures the accessible conformational space under low-salt conditions.

Analysis of methyl relaxation across hydrophobic clusters (Cl1–Cl4) shows that fast side-chain dynamics remain highly consistent across inactive (ensemble 4) and active (ensemble **14**) conformations. Clusters within the PCASP β-sheet (Cl3–Cl4) and the Ig3 domain (Cl1) are minimally affected by Loop 3 rearrangements, indicating that these hydrophobic groups act as structural anchors. Even the interface cluster Cl2, positioned near the active site, displays nearly identical relaxation behaviour in both ensembles. These results demonstrate that the global conformational transitions of the PCASP domain have little influence on fast methyl dynamics, underscoring the stabilizing role of hydrophobic clusters within the MALT1(PCASP–Ig3) scaffold.

In principle extending trajectory analysis to the microsecond regime enables access to motions beyond the temporal range of NMR. This represents an additional strength of our approach: once an MD trajectory segment is selected, it already contains information about conformational motions that remain experimentally inaccessible.

## 4.0 Conclusion

Our results show that under low-salt conditions, monomeric MALT1(PCASP–Ig3)₃₃₉–₇₁₉ consistently collapses into a unified inactive-like ensemble, driven by inward rotation of W580 and coordinated rearrangements of Loops 2 and 3. This reproducible convergence indicates that the inactive conformation is the thermodynamically preferred state of the monomer in the absence of ligand or dimerization.

Changes in ionic strength markedly reshape conformational free-energy landscape of MALT1(PCASP–Ig3)₃₃₉–₇₁₉. Physiological and kosmotropic conditions partially restore access to active-like loop arrangements that remain inaccessible at low salt, whereas high-salt buffers rigidify the PCASP–Ig3 module, suppress loop mobility, and prevent transitions between inactive and active states, thereby strongly stabilizing the active conformation.

MD ensembles initiated from the AF/NMR-derived structure most accurately reproduce backbone dynamics and align closely with ensembles generated from the inactive X-ray model (3V55). This consistency demonstrates that the NMR-derived inactive conformation provides a robust and physically meaningful starting point for characterizing monomeric MALT1.

Backbone-dynamics analysis further reveals that while the structural core of MALT1(PCASP–Ig3)₃₃₉–₇₁₉ is highly stable, Loops 2 and 3 display pronounced, conformation-dependent flexibility. These regions exhibit slow, long-timescale motions that underpin transitions between inactive and active states and ultimately define the protein’s conformational plasticity.

Together, these findings establish that an integrated AF–MD–NMR strategy provides a reliable framework for mapping the intrinsic conformational landscape of monomeric, ligand-free MALT1 and offers a powerful means of resolving regulatory motions that remain inaccessible to experiment alone.

## 5.0 Method

### 5.1.1 Expression of isotope-labelled MALT1(PCASP-Ig3)_339–719_ and preparation of NMR samples

Expression, purification and NMR sample preparation was fully explained in^41^. Shortly the DNA sequence encoding for the PCASP and Ig3 domains of human MALT1, corresponding to residues 338–719 and a C-terminal His6-tag was cloned into the expression vector pET21b (Novagen). MALT1(PCASP-Ig3)_339–719_ was transformed into *Escherichia coli* strain T7 express competent cells (NEB) and expressed in different isotopic labelling combinations in^1^/^2^H,^15^N,^12^/^13^C-labelled M9 medium. One hour prior to induction, precursors were added to the growth medium as previously described^63^. For precursors, 70 mg/L alpha-ketobutyric acid, sodium salt (^13^C4, 98%, 3,3-^2^H, 98%) and 120 mg/L alpha-ketoisovaleric acid, sodium salt (1,2,3,4-^13^C4,99%, 3, 4, 4, 4, –^2^H 97%) (CIL, Andover, MA) were used. Cells were harvested and lysed using ultra-sonicator followed by centrifugation to remove cell debris. The supernatant containing MALT1 was purified by Ni^2+^ Sepharose 6 Fast Flow (Cytiva). A Q-Sepharose HP column (Cytiva) was used to separate monomeric MALT1(PCASP-Ig3)_339–719_ from the dimer form. A final size exclusion chromatography (SEC) step was performed using a HiLoad 16/600 Superdex 200 prep grade column (Cytiva), with running buffer 20mM HEPES 7.4, 50mM NaCl, 1mM DTT. The final monomer MALT1(PCASP-Ig3)_339–719_ protein sample was subsequently exchanged to a buffer (10 mM Tris 7.6, 50mM NaCl, 2mM TCEP, 0.002% NaN_3_, 10% D_2_O) using gravity flow PD10 desalting column (Cytiva). The purified monomeric MALT1(PCASP-Ig3)_339–719_-His protein was concentrated to at least 0.3-0.5 mM for NMR data acquisition.

### 5.1.2 Assessment of the Oligomeric State of MALT1(PCASP-Ig3)_339-719_ at High Salt Concentration

The oligomeric state of the MALT1(PCASP-Ig3)_339–719_ construct was evaluated under high-salt conditions (0.5 M NaCl) prior to MD simulations. Previous LS-MS and chromatography studies have shown that this construct is predominantly monomeric in solution, with dimerization observed only upon binding to peptidic ligands^12, 13^. Some reports also describe mixtures of monomers and dimers in the apo state after purification^22^. To confirm the oligomeric state in our experimental conditions, uniformly ¹⁵N-labeled MALT1 was analysed by gel filtration at 0.5 M NaCl. A single, well-defined peak corresponding to the monomeric species was obtained, and the sample remained stable for at least 24 h (**Figure S7a,b**). To assess whether ionic strength alters the conformational ensemble, TROSY-based ¹H-¹⁵N HSQC spectra were recorded for ¹⁵N-labeled MALT1 at low (0.06 M) and high (0.5 M) NaCl. The overlaid spectra (**Figure S8a**) show near-identical peak patterns, indicating no major structural rearrangements between conditions. Only small chemical shift perturbations were detected, consistent with localized surface electrostatic effects.

As both gel filtration and HSQC measurements confirmed a stable monomeric state of MALT1(PCASP-Ig3)₃₃₉–₇₁₉ at 0.5 M NaCl, MD simulations were subsequently performed under high-salt conditions to investigate the influence of ionic strength on monomer conformational dynamics.

### 5.2 NMR relaxation experiments and data processing

All NMR data were acquired at 298 K on 800 and 900 MHz Bruker spectrometers equipped with TCI Cryo probes which are optimised for triple resonance experiments on biological macromolecules. All spectra were processed using either the mddnmr^64^ and the NMRPipe^65^ softwares at the NMRbox server (https://nmrbox.nmrhub.org/)^66^ or with the TopSpin 4.3.0 (Bruker, Billerica, MA, USA) and analysed using CcpNmr2.4.2^67^ and Dynamics Center 2.8.4 (Bruker, Billerica, MA, USA). Molecular graphics were prepared with the use of Chimera 1.16 and ChemeraX^68, 69^

#### 5.2.1 Determination of^1^H-^15^N CSA/DD cross correlation (η_xy_) relaxation

A complete set of^1^H-^15^N CSA/DD cross correlation relaxation rates, η_xy_, for the backbone amides were acquired at 900 MHz using pulse sequence described in^43^. Experiments were performed using NS=40 on a time domain grid of 1 K x 100 complex points with spectral width/acquisition time of 16 ppm/71 ms for^1^H and 40 ppm/27 ms for^15^N dimensions with D1 = 1s and at a constant-time delay of T = 0.03s. η_xy_ values were determined from a series of 8 relaxation delays: 0.000, 0.002, 0.005, 0.009, 0.014, 0.023, 0.030, 0.044. Carrier positions:^1^H, H_2_O frequency (4.7 ppm);^13^C, 95 ppm;^15^N, 118.0 ppm. Mirror image linear prediction was used for constant-time^15^N sampling.

#### 5.2.2 Determination of Methyl R_1_ and Cross-Correlation Γ_2_ Relaxation

Relaxation measurements for methyl^13^C^1^H_3_ groups^70, 71^ were performed at 800MHz using an interleave pseudo 3D spectral acquisition. Experimental longitudinal relaxation rates (R_1_) of^13^C^1^H_3_ groups were obtained as previously described^61, 72^ by applying mono-exponential fitting to cross peak intensities across a series of 12 two-dimensional correlation spectra. These spectra were recorded with T1 relaxation delays of 0.01, 0.04, 0.08, 0.13, 0.20, 0.29, 0.41, 0.57, 0.69, 0.99, 1.39 to 2.00 s. The number of transients was set to 16.

The dipolar CH, CH cross-correlation contribution to R_2_ (named in this study as Γ_2_) for^13^C^1^H_3_ groups was measured as previously described^71, 72^, using a constant time period of 28.6ms and 14 evolution delays (Δ) of 0.01, 0.6, 1.2, 1.8, 2.4, 3.0, 3.6, 4.2, 4.8, 5.6, 6.4, 7.2, 8.0 to 9.2ms. The number of transients was set to 32.

All parameters in experiments for measuring R_1_ and Γ_2_ relaxation on^13^C^1^H_3_ groups, were set as previously described^61, 72^. The^1^H and^13^C carrier frequencies were set to the water resonance at 4.7 and 16.5 ppm, respectively. The spectral width (SW) for ¹H was 12 ppm over 1024 complex points, while for ¹³C it was 16 ppm over 80 complex points. The inter-scan delay was set to 1 s.

Processing of R_1_ and Γ_2_ spectral datasets was performed using TopSpin4.3.0 software (Bruker) and analysed with the Mathematica software package (Wolfram Research Inc.) as previously described^61, 72^.

### 5.3 Molecular Dynamic simulation

#### 5.3.1 The Protocol used in MD Simulations

This study investigates intramolecular electrostatic interactions and their impact on the time-resolved conformational dynamics of MALT1 monomers, referred to as “4D structural biology” to denote the time-dependent three-dimensional ensembles obtained from molecular dynamics (MD) simulations^73^.

A well-recognized limitation of non-polarizable all-atom force fields is the accurate representation of charged groups^74^. To address this issue, several studies have proposed charge scaling schemes for both proteins and ions^75–77^. Although the TIP3P water model remains the *de facto* standard due to its broad compatibility with biomolecular force fields, it has been argued to insufficiently capture the complexity of water–ion and water–protein electrostatic interactions^78, 79^. Accordingly, careful consideration of simulation protocols was a prerequisite for this study.

Because intramolecular electrostatic interactions are most pronounced under conditions of low ionic strength, owing to reduced electrostatic shielding, we systematically evaluated and benchmarked alternative MD protocols against experimental NMR relaxation data collected under low-salt conditions.

Two molecular dynamics protocols were systematically evaluated: (1) CHARMM19 force field with CUFIX corrections and (2) polarizable AMOEBA model, which explicitly accounts for polarization effects in proteins, ions, and, most critically, water molecules.

#### 5.3.2 MD Simulations Using the Classical Non-Polarizable CHARMM36 Force Field

MD simulations were performed using GROMACS version 2023.1^80^ with the all-atom force field charmm36-mar2019_cufix.ff^81–83^, including a refinement of Lennard-Jones parameters (CUFIX)^84^. Both the protein and the TIP3P water model were treated according to recommended parameters. The initial structures were based on MALT1(PCASP-Ig3)_339–719_ sequence, as detailed in **Table 1**.

The protein was placed at the centre of a periodic cubic box (104 Å), with 55 Na^+^ and 41 Cl^−^ions added to match the 60 mM ionic strength or 415 Na^+^ and 401 Cl^−^ ions added to match the 500 mM ionic strength and maintain electro-neutrality, given the protein’s total charge of –14. For MD with trisodium citrate (3Na^+^ and C_6_H_5_O_7_³⁻) ions, an additional parameterization of citrate ions with charge –3 was performed in the CGenFF software package^85, 86^. To achieve target sodium ion (Na⁺) concentrations of 500 mM and 1500 mM, corresponding amounts of 416 and 1355 Na⁺ ions, along with 134 and 447 citrate (C₆H₅O₇³⁻) ions, were added respectively. Since residues charges were calculated according to pH 7.6, all histidine residues remained neutral. A cutoff of 12 Å was applied for both long-range electrostatic and van der Waals interactions.

To ensure a well-defined starting structure, energy minimization was performed, achieving convergence at a maximum force below 1000 kJ/mol/nm per atom.

Equilibration of the system was performed at 298 K through two restrained phases. Each phase lasted 100 ps and employed positional restraints on protein’s heavy atoms. The initial phase used the NVT ensemble (constant number of particles, volume, and temperature) to stabilize the system temperature. The second phase used the NPT ensemble (constant number of particles, pressure, and temperature) to allow the system density to adjust. Following this, all restraints were removed, and a free production molecular dynamics simulation was carried out in the NPT ensemble for 2500 ns until the system reached stability, indicated by a plateau in root mean square deviation (RMSD) values.

A modified Berendsen-type (V-rescale) thermostat and a Parrinello-Rahman barostat were employed. Hydrogen-containing covalent bonds were constrained using the Links algorithm with a 2 fs timesteps. Following equilibration, MD simulations were continued as a production run for 500 ns under the same conditions. System stability was assessed using standard GROMACS tools^80^ including control of the temperature, pressure, energy, secondary structure, the box border, and RMSD.

#### 5.3.3 Molecular Dynamics Simulations Using the Polarizable AMOEBA Force Field

Electrostatic interactions play a crucial role at low ionic strength due to reduced shielding. Therefore, an MD simulation was performed in Tinker-9 using the polarizable AMOEBA force field to assess whether the classical non-polarizable force field sufficiently agrees or not with experimental results.

The protein residue charges were calculated at pH 7.6, in the same way as for GROMACS in the previous section, with all histidine residues remaining neutral. The ionic strength was set to 60 mM, accounting for both buffer and salt concentrations.

MD simulations were performed using Tinker-9 [Tinker9: Next Generation of Tinker with GPU Support. Zhi Wang, Jay W. Ponder, 2021, https://github.com/TinkerTools/tinker9] with the all-atom polarizable force field AMOEBABIO18^87^. The protein was placed at the centre of a periodic cubic box (103.9375 Å), with 55 Na^+^ and 41 Cl^-^ ions added to match the desired ionic strength and maintain electro-neutrality, given the protein’s total charge of –14. Molecular dynamics simulations were performed using the RESPA integrator with a 2 fs outer time step and a preconditioned conjugate gradient polarization solver (with a 10^-5^ convergence threshold). Periodic boundary conditions (PBCs) were applied using the Smooth Particle Mesh Ewald (SPME) method, with a 120 × 120 × 120 Å grid. The Ewald-cut off was set to 7 Å, while van der Waals and electrostatic charge-charge cut offs were 12 Å. Mutual dipole polarization was applied to iterate induced dipoles to self-consistency, with a convergence cutoff of 10^−5^ Debye. Before the simulation, the protein structure underwent energy minimization to optimize starting structure geometry and solvent orientation. Convergence was achieved at a maximum force below 0.01 kcal/mol/Å per atom.

Equilibration was performed in two phases. First, NVT ensemble was applied 100 ps equilibration. The system was heated to 298 K until the temperature plateaued. Second, NPT ensemble 100 ps was used for the continued equilibration until pressure and density stabilized. A Bussi-Parrinello stochastic thermostat and a Monte Carlo barostat were used. The RESPA integrator was applied. Following equilibration, MD simulations continued as a production run for 3000 ns under the same conditions. System stability was monitored using standard Tinker9 tools, tracking temperature, pressure, energy and periodicity.

#### 5.3.4 Starting structures for MD-simulations

Both the experimental X-ray structures (PDB 3V55) and the predicted structure from AlphaFold [https://www.nature.com/articles/s41586-024-07487-w] were used as starting structures for the MALT1 protein in solution. Structural modifications and modelling of the missing loops regions (pdb id 3v55) and missing α-helix region (pdb id 9mkd) of the crystal structure were performed with PDBFixer^88^. However, the primary focus of this work was not only the initial conformations of MALT1 but also the solution properties, such as ionic strength (IS), while pH 7.6 and temperature of 298 K remained constant for all MD runs. The combinations of starting conformations (SCs) of MALT1 and ions concentrations used in the simulations are presented in **Table 1**.

### 5.4 MD Trajectory Analysis: Alignment, RMSD Calculation, and Back-Calculation of NMR Relaxation Parameters

Structural alignments and subsequent RMSD calculations were performed using two distinct sets of backbone heavy atoms 1: [342-717] and 2: [342-467, 484-491, 509-562, 572-717], excluding the mobile N– and C-terminal residues (338-341and 718-725) in both cases. In the second set structures were aligned excluding the mobile inter-domain linker (563-571) and flexible loops (468-483 and 492-508), followed by an RMSD calculation for both sets of backbone heavy atoms. All RMSD values were computed using the GROMACS software package.

Principal component analysis (PCA) was performed on Cartesian coordinates for all molecular dynamics (MD) trajectories using the GROMACS software package. The analysis was conducted on the backbone heavy-atoms, analogous to those used for the RMSD calculation. For each trajectory, 30 eigenvalues were calculated. The data for the first three eigenvalues corresponding to the first three principal components (PCi) are presented in **Table 2**.

**Table 2.** The first three eigenvalues for the first three principal components (PCs) for trajectories.

| № MD trajectory | Backbone heavy-atoms of stable regions 1<sup>a</sup> |  |  | Backbone heavy-atoms of stable regions 2<sup>b</sup> |  |  |
| --- | --- | --- | --- | --- | --- | --- |
|  | PC1<sup>c</sup> | PC2 | PC3 | PC1 | PC2 | PC3 |
| 3 | 7.4 | 3.9 | 2.5 | 3.6 | 1.8 | 1.2 |
| 4 | 19.3 | 3.2 | 2.3 | 7.5 | 1.6 | 1.1 |
| 5 | 12.7 | 6.9 | 4.4 | 6.7 | 4.0 | 2.8 |
| 6 | 103.1 | 59.7 | 28.5 | 65.1 | 51.7 | 20.8 |
| 8 | 4.9 | 3.5 | 1.6 | 3.0 | 2.8 | 1.4 |
| 9 | 8.2 | 2.9 | 1.7 | 2,5 | 1,4 | 0,8 |
| 11 | 20.9 | 7.7 | 5.2 | 4.1 | 2.0 | 1.7 |
| 14 | 20.7 | 3.2 | 2.6 | 3.6 | 1.9 | 1.1 |
| 15 | 32,8 | 4,8 | 3,5 | 21,4 | 2,4 | 1,7 |
| 16 | 11.2 | 8.2 | 5.9 | 5.5 | 3.6 | 2.3 |

<sup>a</sup> region used to perform PCA: 342-717
<sup>b</sup> regions used to perform PCA: 342-467, 484-491, 509-562, 572-717
<sup>c</sup> λ eigenvalue for each PC<sub>i</sub> (nm<sup>2</sup>)

RMSF values were calculated for Cα atoms after centre-of-mass alignment of all trajectory frames and was computed using the standard definition^89^.

### 5.5 Individual MD trajectory analyses with back-calculation of theoretical^15^N and^13^C relaxation parameters

The back-calculation of NMR relaxation parameters followed the method described in our earlier publications^43, 61^, using the final 500 ns of the trajectory for calculating the correlation function with maximal time of 7×τ_c_, with an experimental overall correlation tumbling of τ_c_ =27ns. MD trajectory regions were analysed by back-calculation NMR spin-relaxation parameters, using a bootstrapping procedure to estimate parameters dispersion, as previously described^61^. Each 500 ns MD segment was selected to be several times longer than the maximum duration of the autocorrelation function acf(t), specifically exceeding 7×τ_c_, to ensure proper averaging of acf(t) values and effective application of the moving block bootstrap method^61^. For each MD segment, the analysis began by aligning all protein frames to the mean structure, using the heavy atoms of rigid backbone residues.

The backbone^1^H-^15^N vector extraction and approximation of autocorrelation function acf(t) to a multi-exponential decay

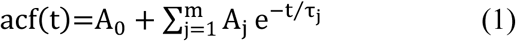

with the best-fit parameters A_0,_ A_j,_ τ_j_ and the subsequent spectral density function J(ω) calculations were utilized as previously described^43, 61^, using homebuilt scripts in “Mathematica” software package [Wolfram Research] and the MD Analysis external library [mdanalysis.org]. The resulting acf(t) values could be calculated with respect to t values (0.1ns, 1ns, 10ns, 100ns) presenting the time scale of the motions (Figure 8).

**Figure 8.**
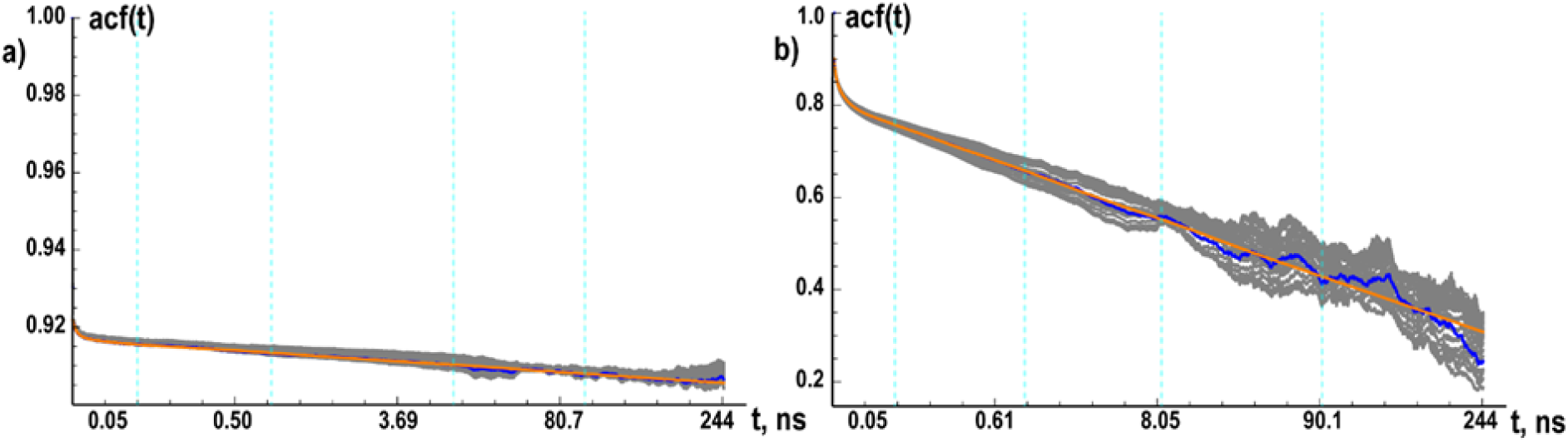
Examples of backbone NH autocorrelation decay for residues 365D and 571A. Representative backbone NH autocorrelation functions (acf) illustrating the decay over time (ns) are shown for residues 365D (a) and 571A (b). The fitted approximation, residue-averaged values, and deviations from the MD simulations are shown in orange, blue, and grey, respectively. Blue dashed lines indicate the acf(t) values at 0.1 ns, 1 ns, 10 ns, and 100 ns for the relatively stable NH vector of 365D (a) and the more flexible NH vector of 571A (b).

Back-calculation of classical NMR^15^N relaxation parameters η_xy_, R_1,_ R_2_ and NOE as a function of J(ω) were also performed as following:

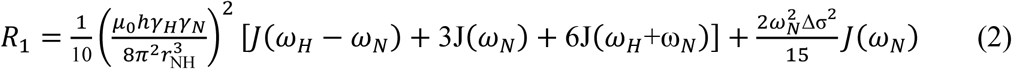

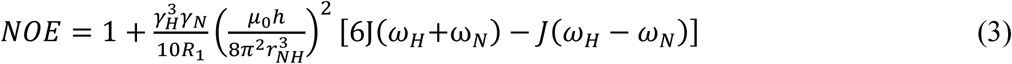

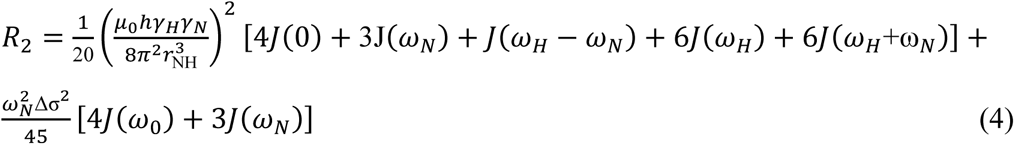

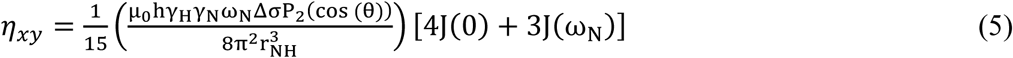

Where the spectral density function was:

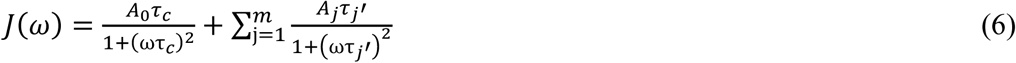

Where 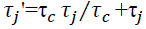 and τ_*c*_ is the experimental rotation correlation time, µ_*o*_ is the vacuum permeability; h is Planck’s constant; *Y*_*H*_ and *Y*_*N*_ are the gyromagnetic ratios of^1^H and^15^N respectively; Δσ is the chemical shift anisotropy (CSA) of^15^N with Δσ = −166±9ppm^90^; r_NH_ = 1.023±0.006 Å^91^; ω_N_ and ω_H_ are the Larmor frequencies of^15^N and^1^H at 800 or 900 MHz, respectively; CSA tensor value with respect to the NH vector ΔσP_2_(Cos(θ)) = −145±8ppm, θ is the CSA/NH vector angle and P_2_ is the Legendre 2nd degree polynomial^92^; J(ω) is the NH auto-correlation spectral density function.

### 5.6 Identifying Stable Conformational Ensembles from Molecular Dynamics Simulations

To identify regions within the trajectory where the protein samples stable conformational ensembles (persisting longer than 10 correlation times, τ_c_), we employed the following method. The approach analyses the populations of clusters identified via an RMSD-based criterion. First, we performed cluster analysis (Gromos algorithm, RMSD cutoff of 0.1 nm) on a molecular dynamics production trajectory with a 1 ns frame interval. This assigns every trajectory frame to a specific cluster. The trajectory was then divided into overlapping time windows (15 ns step, 10*τ_c_ length). For each window, the population of each cluster was calculated (Figure 9).

**Figure 9.**
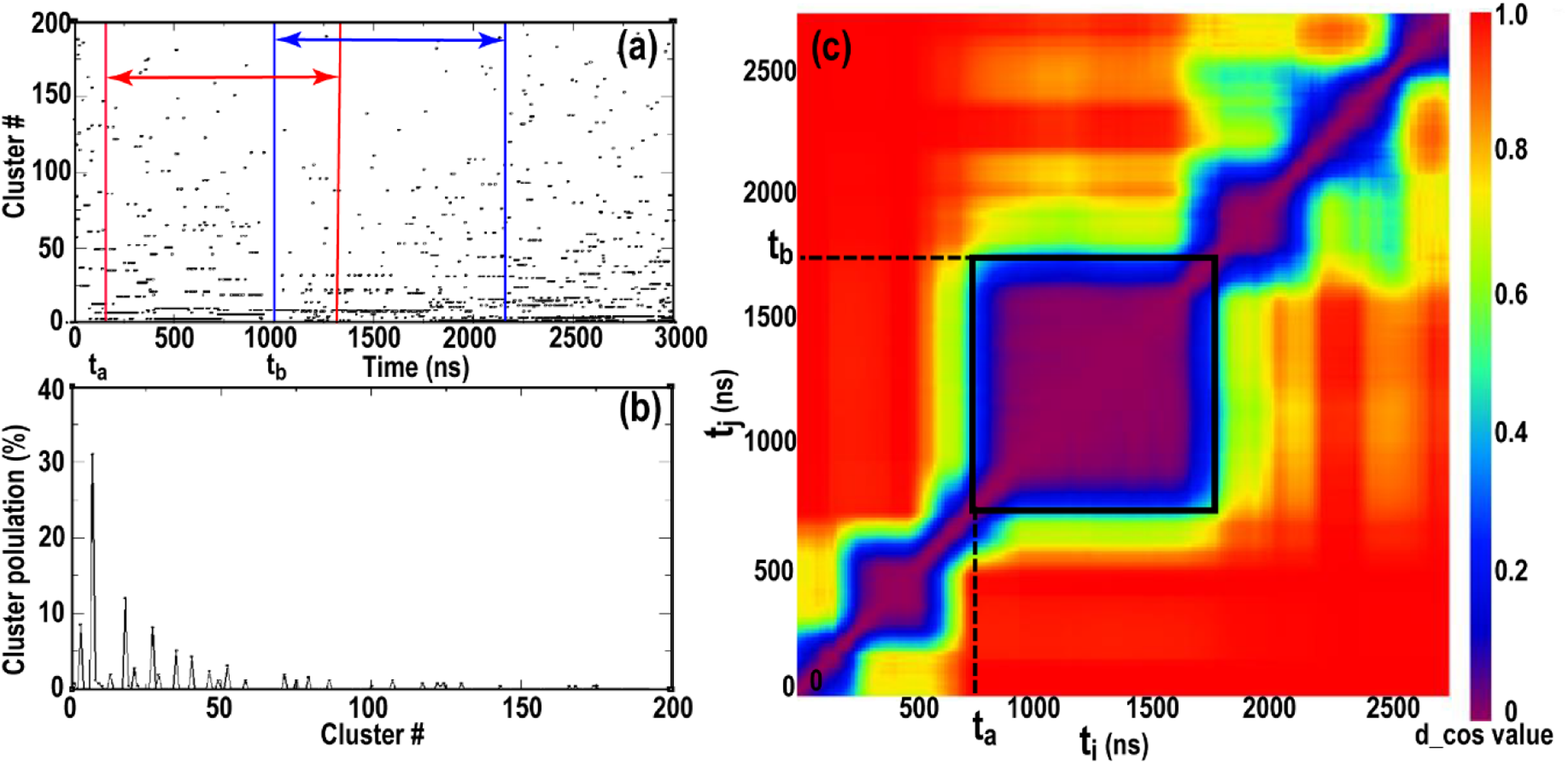
Cluster analysis and stability evaluation of the 4th MD trajectory. (a) Distribution of frames from the 4th MD trajectory (sampled every 1 ns) across structural clusters. Clustering was performed using an RMSD cutoff of 0.1 nm, such that the pairwise RMSD between any two frames within a cluster is ≤0.1 nm. Overlapping time windows of duration 10*τ_c_, initiated at t_i_ = t_a_ and t_j_ = t_b_, are indicated by red and blue lines, respectively. (b) Cluster populations (%) for the time window [t_a_; t_a_+10*τ_c_]. Each time window is represented by a vector 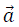, whose components correspond to the percentage populations of the individual clusters. (c) Cosine distances calculated for all pairs of cluster-population vectors derived from the 4th trajectory. Results are shown as a heat map. The region of temporal stability used for subsequent analysis is outlined by a black square, beginning at t_a_ (750 ns), ending at t_b_ (1785 ns), and spanning a duration of t_b_ + 10*τ_c_ – t_a_=1292 (ns).

For a time window with an initial time t_i_ (where t_i_ = tₐ represents a specific instance of t_i_), spanning from tₐ to tₐ + 10*τ_c_, the population of each cluster is quantified. The frames in the time window, each assigned to a specific cluster. This allows for the computation of the percentage of frames belonging to each cluster, which is formalized as a population vector 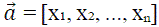. Where, the vector index corresponds to a distinct, sequentially numbered cluster (including clusters with population equal 0), and its value xₖ denotes the percentage of total frames attributed to the k-th cluster. The total number of frames in this analysis is fixed at 10*τ_c_*(frame interval). Analogously, for a distinct time window with initial time tⱼ = t_b_, the cluster populations are given by a corresponding vector 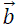.

The similarity between the two vectors was quantified using the cosine distance^93^:

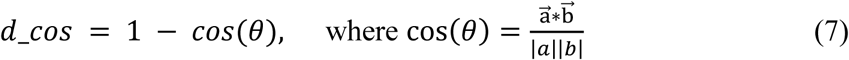

where vectors 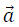 and 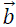 represent the cluster populations for windows starting at times t_a_ and t_b_, respectively. A value of 0 indicates identical distributions, while 1 denotes complete dissimilarity. The resulting pairwise cosine distance matrix was visualized as a heatmap.

A stable conformational ensemble was defined by the presence of a square block along the diagonal of this matrix, bounded by times t_a_ and t_b_, where all elements were below a threshold of 0.1^94^. The ensemble corresponding to this block is characterized by the time interval from t_a_ to t_b_. However, since the analysis is performed by partitioning the trajectory into time windows of duration 10*τ_c_, the boundary point t_b_ also defines such a window, starting at t_b_ and ending at t_b_ + 10*τ_c_. Consequently, the final trajectory segment describing the stable conformational ensemble is defined by the interval from t_a_ to t_b_+10*τ_c_.

### 5.7 Penalty functions for MD trajectory validation

An approach was used to identify the most representative molecular dynamics (MD) trajectory that agrees well with experimental NMR relaxation data. The ranking was performed by comparing back-calculated relaxation parameters (η_xy_, NOE, R₁, and R₂) back-calculated from the MD trajectories with the corresponding experimental one. First, both theoretical and experimental values were normalized with respect to the corresponding mean experimental values:

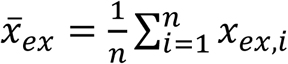

Where I is residue number, 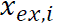 is i-th experimental value of the relaxation parameter, 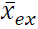 is the average value of the relaxation parameter and n is the number of values used in the ranking.

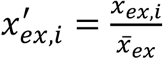

where 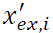 is the new normalized experimental i-th value.

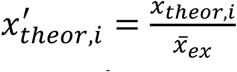

where 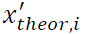, n is back-calculated i-th value of the relaxation parameter for MD trajectory and 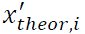 is the new normalized i-th value.

The analysis included data from amino acid residues, excluding the following regions: 340–350, 364–400, 407–419, 444–448, 455–462, 487–488, 525–538, 578–621, 631–639, 644–655, 679–692, 700–715, and the His-tag. Based on these data, the ranking was conducted according to three principal metrics: mean absolute error (MAE), root mean square error (RMSE)^95^ and cosine distance^93^.

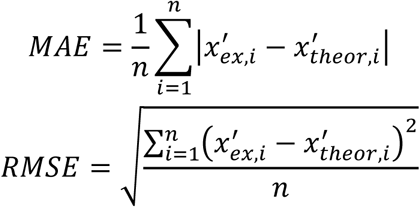

For the calculation of the cosine distance, the experimental parameter values were arranged into a vector 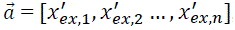, while the theoretical values were arranged into a vector 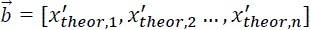 The cosine distance was computed using the equation (7). The data of penalty functions are presented in **Table 3**.

**Table 3.** Penalty functions.

| # tr<sup>a</sup> /RP<sup>b</sup> | R1 (900MHz) | R2 (900MHz) | ($\eta_{xy}$ (900MHz) | R1 (800MHz) | NOE (800MHz) |
| --- | --- | --- | --- | --- | --- |
| 3<sup>c</sup> | 0.2016 | 0.1315 | 0.1416 | 0.2435 | 0.1818 |
| 3<sup>d</sup> | 0.3170 | 0.1756 | 0.1894 | 0.3583 | 0.1894 |
| 3<sup>e</sup> | 0.0396 | 0.0131 | 0.0162 | 0.0468 | 0.0296 |
| 4<sup>c</sup> | 0.1897 | 0.1195 | 0.1396 | 0.2368 | 0.2044 |
| 4<sup>d</sup> | 0.2919 | 0.1659 | 0.1905 | 0.3416 | 0.1905 |
| 4<sup>e</sup> | 0.0321 | 0.0118 | 0.0168 | 0.0400 | 0.0343 |
| 5<sup>c</sup> | 0.2302 | 0.1676 | 0.1802 | 0.2564 | 0.2259 |
| 5<sup>d</sup> | 0.3450 | 0.2196 | 0.2322 | 0.3651 | 0.2322 |
| 5<sup>e</sup> | 0.0536 | 0.0186 | 0.0229 | 0.0576 | 0.0392 |
| 8<sup>c</sup> | 0.2316 | 0.1361 | 0.1619 | 0.2599 | 0.1837 |
| 8<sup>d</sup> | 0.3542 | 0.1735 | 0.2127 | 0.3853 | 0.2127 |
| 8<sup>e</sup> | 0.0449 | 0.0136 | 0.0214 | 0.0499 | 0.0307 |
| 9<sup>c</sup> | 0.2467 | 0.1390 | 0.1700 | 0.2788 | 0.1927 |
| 9<sup>d</sup> | 0.3721 | 0.1871 | 0.2319 | 0.4040 | 0.2319 |
| 9<sup>e</sup> | 0.0494 | 0.0166 | 0.0257 | 0.0565 | 0.0309 |
| 16<sup>c</sup> | 0.2513 | 0.1600 | 0.1865 | 0.2822 | 0.1934 |
| 16<sup>d</sup> | 0.3997 | 0.2191 | 0.2524 | 0.4243 | 0.2524 |
| 16<sup>e</sup> | 0.0689 | 0.0227 | 0.0304 | 0.0740 | 0.0320 |

<sup>a</sup> Number of the MD trajectory (as in the table 1)
<sup>b</sup> NMR Relaxation parameter
<sup>c</sup> Value of the mean average error (MAE)
<sup>d</sup> Value of the root mean square error (RMSE)
<sup>e</sup> Value of the cosine distance (d<sub>cos</sub>)

### 5.8 MD Simulation Protocol for MALT1(PCASP-Ig3)_339–719_ Dynamics at Low Salt Concentration

To generate a conformational ensemble of MALT1(PCASP-Ig3)_339–719_ that accurately reproduces experimental backbone relaxation parameters, we optimized two key aspects of the MD protocol: trajectory length, and force field selection. Since experimental relaxation data are available only under low-salt conditions (20 mM Tris pH 7.5, 50 mM NaCl, 1 mM TCEP), all optimizations were performed in the same buffer environment to ensure meaningful comparison with experiment.

#### 5.8.1 Effect of Trajectory Length on Back-Calculated Relaxation Parameters

Previously, accurate back-calculation of correlation functions act(t) required MD trajectories of sufficient length, typically >7 τ_c_, and extracted from intervals in which the backbone RMSD had reached a stable plateau^43, 61^. In this study, we identified the optimal analysis window by examining multiple intervals within a 6 μs MD simulation using starting structure 4 using RMSD-based clustering to assess conformational populations described above. R₁, R₂, and NOE parameters were evaluated for three segments Figure 1: 1000–3000 ns, 1000–1500 ns, and 2500–3000 ns. Although all segments showed broadly similar agreement with experiment, the 2500–3000 ns interval reproduced the experimental data most closely in the Loop 5 and Loop 6 regions (**Figure S9**), indicating that at least 2.5 μs of equilibration is required for reliable analysis.

#### 5.8.2 Effect of Force Field

Finally, we evaluated whether a polarizable force field improves accuracy under low-ionic-strength (55 Na^+^ and 41 Cl^-^) conditions. Using starting structure **4**, we compared MD simulations performed with the CHARMM36 + CUFIX force field (trajectory **4**) and the polarizable AMOEBA force field (trajectory **5**). Back-calculated R_1_ and η_xy_ relaxation data (Figure **S10**) showed no improvement with AMOEBA. Therefore, the classical CHARMM19 + CUFIX force field provides sufficient accuracy for studying MALT1 conformational dynamics and was adopted for all subsequent simulations.

## AUTHOR CONTRIBUTIONS

XH has contributed with production of all necessary protein variants, including their purifications, and developing a construct of labelled MALT1 proteins. DL, VO and TA wrote the original manuscript draft. TA, AA, TS, AL, VO and PA contributed with final editing of the manuscript. DL contributed to the development of the η_xy_ pulse sequence. V.O. and D.L. contributed with NMR measurements and methodology of MALT1. D.L and KR performed M.Dcalculation and analysis.

## COMPETING INTERESTS

The authors declare no competing interests.

## ACKNOWLEDGEMENTS

The authors thank the Swedish NMR Centre for access to the instruments and support. This work was supported by the Swedish Foundation for Strategic Research grant ITM17-0218 to T.A and P.A.; Swedish Cancer Society (21 1605 Pj 01 H), Cancer och Allergi Fonden (10399), and Swedish Research Council (2021-05061 and 2018-02874) to A.A; RSF 24-13-00413 to D.L. and 2019-03561, 2023-03485 to V.O. This study used NMRbox: National Center for Biomolecular NMRData Processing and Analysis, a Biomedical Technology Research Resource (BTRR), which is supported by NIH grant P41GM111135 (NIGMS).

## DATA AVAILABILITY

The ensembles of AF and MD structures and other information and data are available upon request from the authors. The numerical sources for the graphs and plots are found in the supplementary data. The assignment is found in BiologicalMagneticResonanceDataBank (http://www.bmrb.wisc.edu/) with the BMRB accession code 52265. The data were acquired using TopSpin3.5 and TopSpin4.06 from Bruker.

## DATA AVAILABILITY

The ensembles of AF and MD structures and other information and data are available upon request from the authors. The numerical sources for the graphs and plots are found in the supplementary data. The assignment is found in BiologicalMagneticResonanceDataBank (http://www.bmrb.wisc.edu/) with the BMRB accession code 52265. The data were acquired using TopSpin3.5 and TopSpin4.06 from Bruker. The 4-th trajectory (2500-3000 ns time interval) was clustered into 20 states using an RMSD cutoff of 0.105 nm. The cluster populations were: 1–34.4%, 2–17.7%, 3–13.5%, 4–8.9%, 5–4.4%, 6–3.5%, 7–3.3%, 8–2.7%, 9–1.6%, 10–1.3%, 11–1.0%, 12–1.0%, 13–0.9%, 14–0.7%, 15–0.6%, 16–0.5%, 17–0.5%, 18–0.5%, 19–0.4%, 20–0.3%, and other–2.3%. The most representative 20 structures were selected to construct a conformational ensemble, which was deposited in the Protein Data Bank (PDB ID: D_4-R5HP).

## Notes

### Competing Interest Statement

The authors have declared no competing interest.

