## Supplemental information for "Insight into Malt1 activation mechanism through synergetic approach of AlphaFold, MD Simulation and NMR dynamic analyses"

### Table of Contents

1. Figure S1 Structural superpositions along the PC1 mode under different ionic conditions.
2. Figure S2 | Conformational dynamics of MALT1(PCASP–Ig3)$_{339–719}$ under different ionic conditions. Root-mean-square deviation (RMSD) of individual trajectories by NMR and molecular dynamics. Ensembles of trajectories **8**, **9** and **4**
3. Figure S4 | Backbone amide dynamics of MALT1(PCASP–Ig3)$_{339–719}$ measured by NMR and molecular dynamics. Ensembles of trajectories **3**, **4**, **16**
4. Methyl Group Dynamics as Probes of Conformational Stability: Agreement Between MD and NMR
    a. Figure S5 MALT1(PCASP-Ig3)$_{339–719}$ Ile, Val; Leu Methyl Dynamic Parameters of the $R_1$ and $\Gamma_2$ obtained on 800 MHz spectrometers
5. Relaxation Analysis Across Hydrophobic Clusters
6. Cl2: A Critical Cluster at the Ig3–PCASP Interface
    a. Figure S6. Methyl relaxation dynamics across hydrophobic clusters in MALT1(PCASP-Ig3)$_{339–719}$
7. Assessment of the Oligomeric State of MALT1(PCASP-Ig3)$_{339-719}$ at High Salt Concentration
    a. Figure S7 Salt dependence of the monomeric state of the apo MALT1(PCASP-Ig3)$_{339–719}$ based on Ni-NTA affinity purification
    b. Figure S8 TROSY $^1$H–$^{15}$N HSQC spectra of $^{15}$N-labeled MALT1 at different salt concentrations
8. Figure S9 MALT1(PCASP-Ig3)$_{339–719}$, amide backbone, $^{15}$N(H), dynamic parameters obtained on 800 MHz spectrometers
    a. Figure S10 MALT1(PCASP-Ig3)$_{339–719}$, amide backbone, $^{15}$N(H), dynamic parameters obtained on 900 MHz spectrometers **3**, **4** and **5**

9. Figure S10 MALT1(PCASP-Ig3)$_{339–719}$, amide backbone, $^{15}$N(H), dynamic parameters obtained on 900 MHz spectrometers
10. Figure S11. Conformational ensembles of MALT1(PCASP-Ig3)$_{339–719}$ were obtained from free-restraints MD simulations at low salt concentration. Superposition of 10 structures from trajectory **4**,
11. References

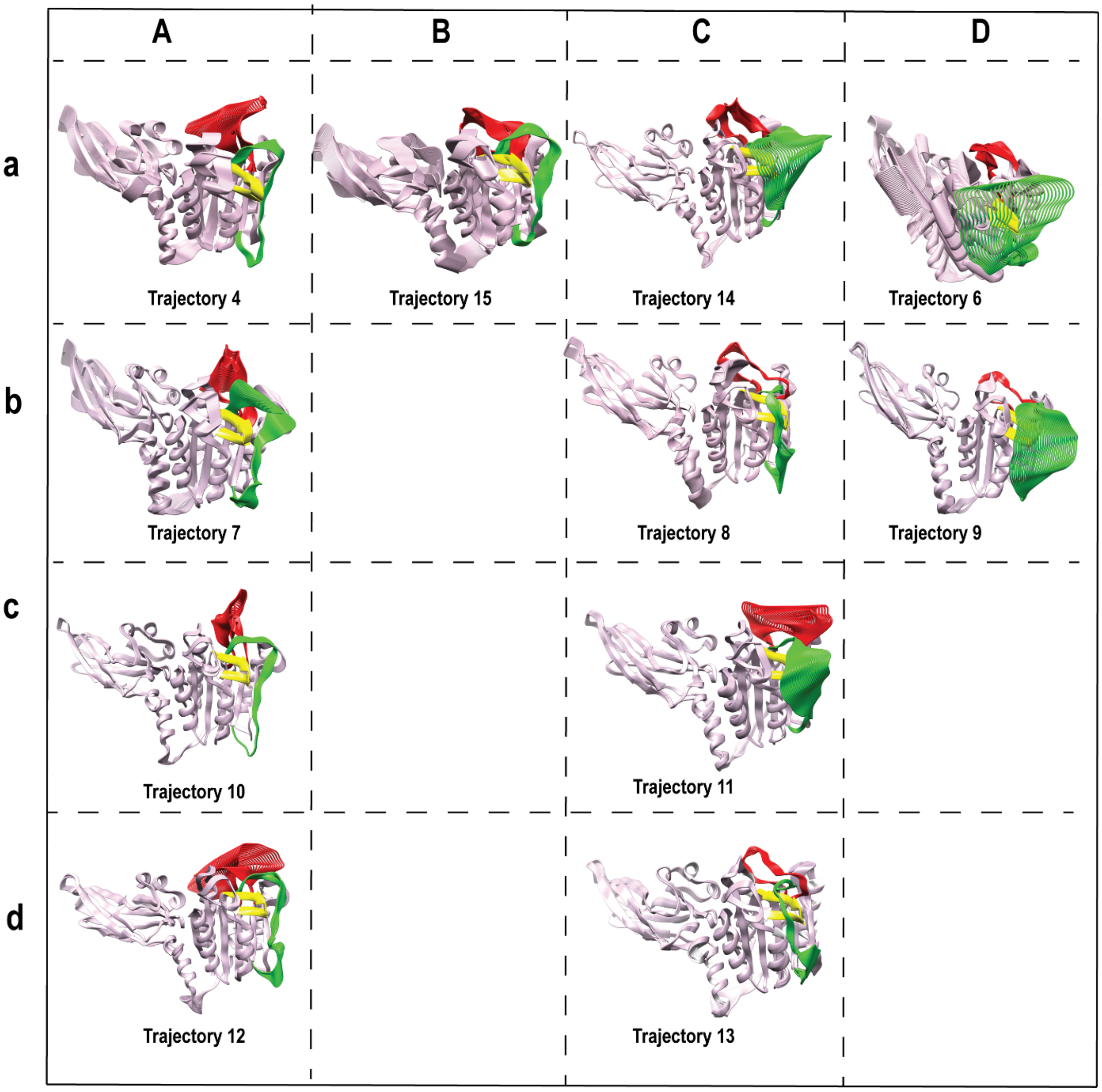


**Figure S1 Structural superpositions along the PC1 mode under different ionic conditions.** Superpositions of 30 structures sampled from opposite ends of the conformational space defined by the first principal component (PC1) for all ensembles. Columns A and B show structures in the inactive conformation, whereas columns C and D show structures in the active conformation. Columns A and C correspond to the W580 **in** conformation, while columns B and D correspond to the W580 **out** conformation. Rows a–d represent, respectively, 60 mM NaCl (55 $Na^+$ 41 $Cl^-$); 500 mM NaCl (415 $Na^+$, 401 $Cl^-$); 166.7 mM sodium citrate (416 $Na^+$, 134 $C_6H_5O_7^{3-}$); and 500 mM sodium citrate (1355 $Na^+$, 447 $C_6H_5O_7^{3-}$).

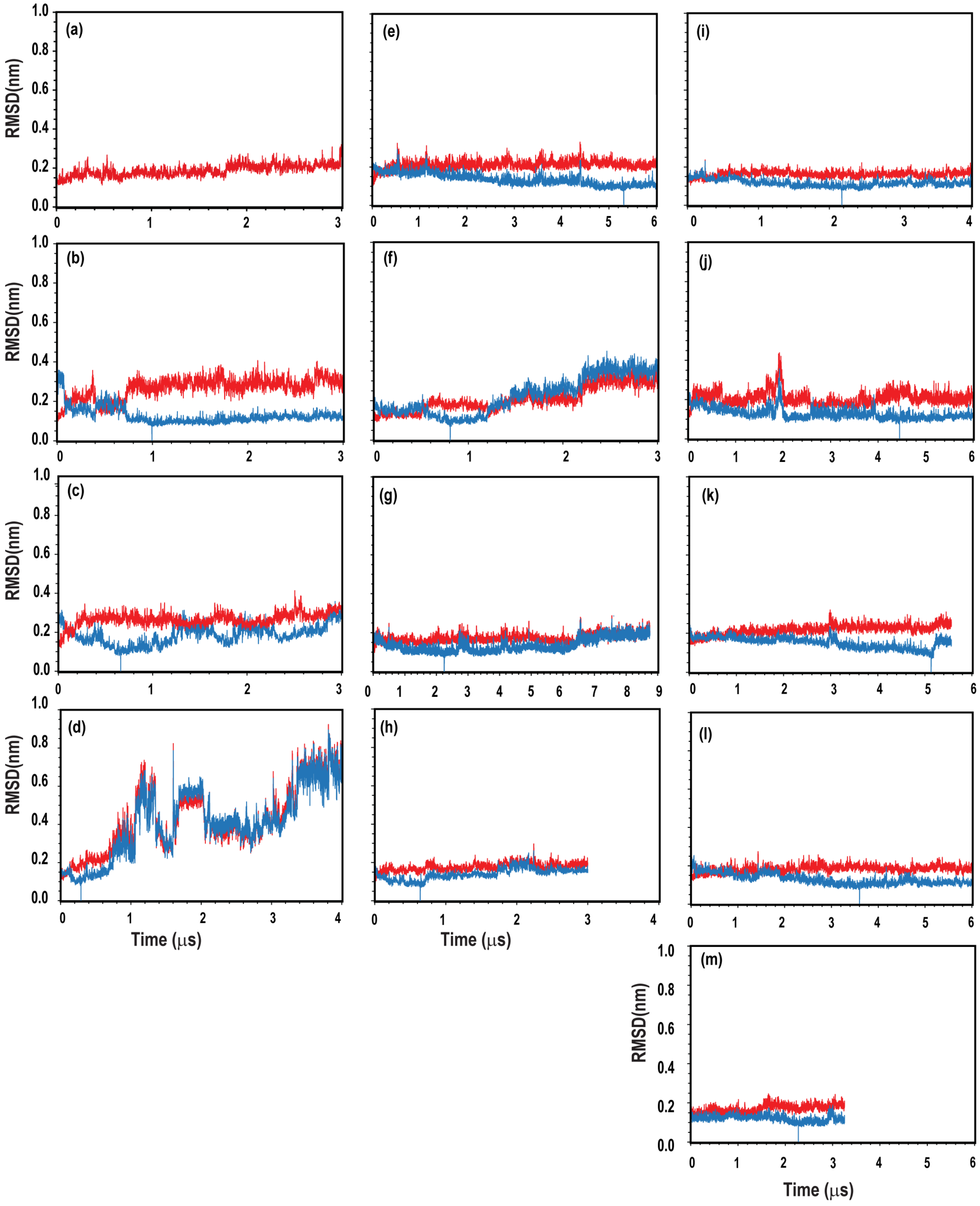


**Figure S2 | Conformational dynamics of MALT1(PCASP–Ig3)$_{339–719}$ under different ionic conditions.** Root-mean-square deviation (RMSD) of individual trajectories relative to their respective starting structures (red) and the best cluster structure (blue) are shown. Panels (a–f) correspond to 60 mM NaCl (55 $Na^+$ 41 $Cl^-$); trajectories 3, 4, 5, 6, 14 and 15, respectively). Panels (g–i) correspond to 500 mM NaCl (415 $Na^+$, 401 $Cl^-$; trajectories 7, 8 and 9, respectively). Panels (j–k) correspond to 166.7 mM sodium citrate (416 $Na^+$, 134 $C_6H_5O_7^{3-}$; trajectories 10 and 11, respectively). Panels (l–m) correspond to 500 mM sodium citrate (1355 $Na^+$, 447 $C_6H_5O_7^{3-}$; trajectories 12 and 13, respectively).

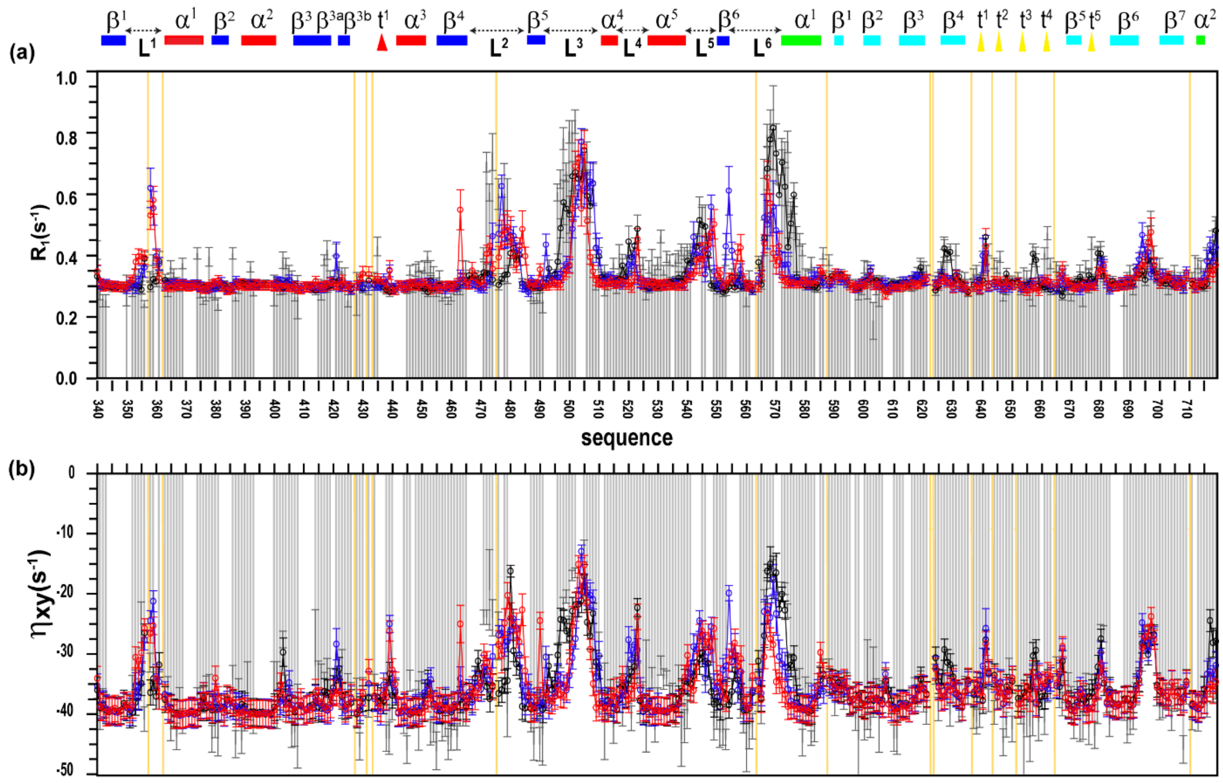

**Figure S3 | Backbone amide dynamics of MALT1(PCASP–Ig3)$_{339–719}$ measured by NMR and molecular dynamics.** Backbone amide relaxation parameters of MALT1(PCASP–Ig3)$_{339–719}$ measured at 900 MHz. Longitudinal relaxation rates $R_1$ ($s^{-1}$) and CSA–dipole cross-correlation relaxation rates ($\eta_{xy}$) are shown for ensembles **8**, **9** and **4**. Experimentally measured $R_1$ and $\eta_{xy}$ values are shown as light-grey bars. Corresponding parameters predicted from molecular dynamics ensembles **8**, **9** and **4** are shown as solid blue, red and black lines, respectively. Ensembles **8** and **9** were generated from trajectory segments 2500–3000 ns and 3500–4000 ns, respectively, at high salt concentration (500 mM NaCl; 415 $Na^+$, 401 $Cl^-$) using the CHARMM36-19 force field. Ensemble **4** was generated from trajectory segments 2500–3000 ns at low salt concentration (60 mM NaCl; 55 $Na^+$, 41 $Cl^-$) using the same force field. Experimental error bars represent one standard deviation from curve fitting, and uncertainties in predicted parameters were estimated by bootstrap analysis.

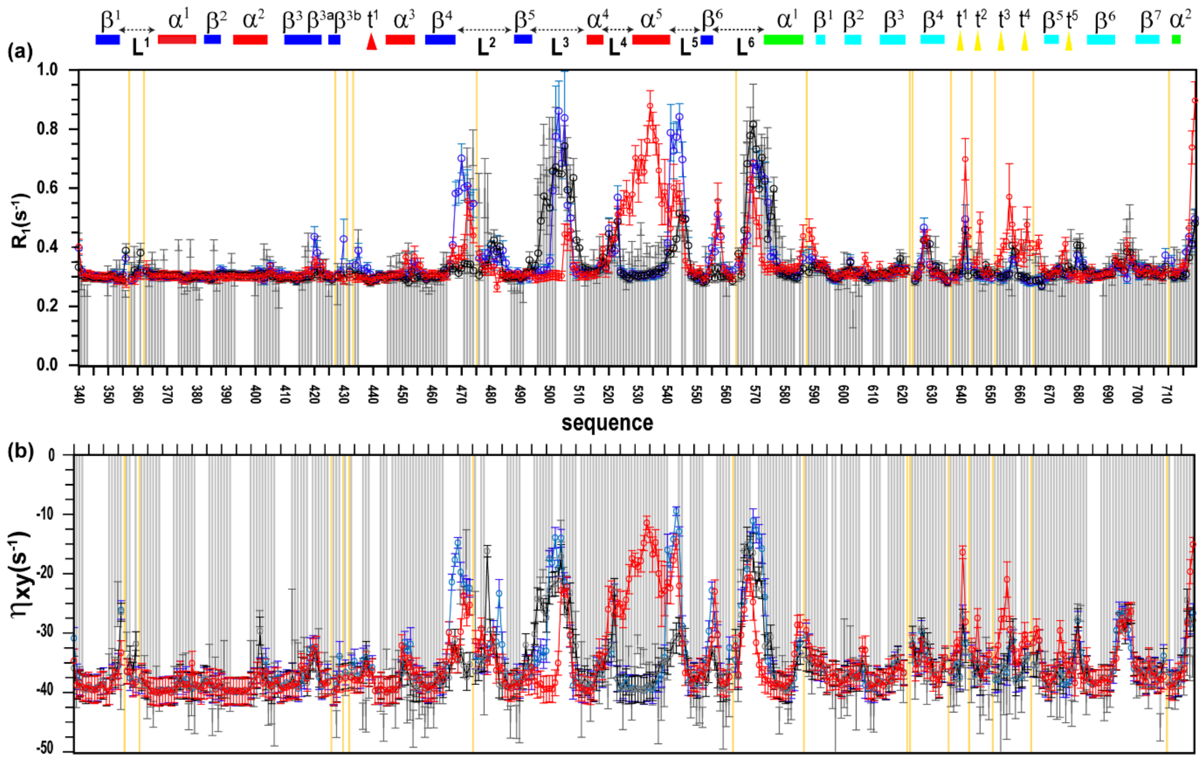

**Figure S4 | Backbone amide dynamics of MALT1(PCASP–Ig3)$_{339–719}$ measured by NMR and molecular dynamics.** Backbone amide relaxation parameters of MALT1(PCASP–Ig3)$_{339–719}$ measured at 900 MHz. Longitudinal relaxation rates $R_1$ ($s^{-1}$) and CSA–dipole cross-correlation relaxation rates ($\eta_{xy}$) are shown for ensembles **3**, **4** and **16**. Experimentally measured $R_1$ and $\eta_{xy}$ values are shown as light-grey bars. Corresponding parameters predicted from molecular dynamics ensembles **3**, **4** and **16** are shown as solid blue, black and red lines, respectively. Ensembles **3**, **4** and **16** were generated from trajectory segments 2500–3000 ns, at low salt concentration (60 mM NaCl; 55 $Na^+$, 41 $Cl^-$) using the CHARMM36-19 force field. Experimental error bars represent one standard deviation from curve fitting, and uncertainties in predicted parameters were estimated by bootstrap analysis.

### 4 Methyl Group Dynamics as Probes of Conformational Stability: Agreement Between MD and NMR

In our previous study on the apo form of human MALT1(PCASP-Ig3)$_{338–719}$ (Han *et al.*, 2022) we reported nearly complete $^{1}H$ /$^{13}C$ methyl resonance assignments for Ile, Leu, and Val residues. These findings provide a solid foundation for further investigation of both backbone and side-chain dynamics in the monomeric form of MALT1.

The longitudinal relaxation rate ($R_1$) and cross-correlated relaxation rate ($\Gamma_2$), both measured at 800 MHz, are shown in light blue solid brackets in **Figure S5**. For methyl groups, $R_1$ values span ~1.0–5.0 $s^{-1}$, while $\Gamma_2$ values range from ~5.0–25.0 $s^{-1}$, reflecting a broad distribution of internal correlation times ($\tau_e$) and order parameters ($S^2$).

Additionally, the $^{13}C$ relaxation parameters for $^{13}C_{d2}$ for methyl group of Leu amino acid should be interpreted with caution due to the possible formation of so-called AB spin systems with $^{13}C_g$, which can occur when the chemical shift difference is smaller than ~350 Hz (≈10 × $^{1}J(C_{d2}C_g)$).

Closer inspection of $R_1$ relaxation reveals clear amino acid–specific differences. Ile methyl groups display the narrowest distribution, with values clustered around 1.0 $s^{-1}$ (**Figure S5e**), despite being distributed across diverse structural elements of MALT1(PCASP-Ig3)$_{338–719}$. By contrast, Val and Leu methyl groups show higher average $R_1$ values (~2.5 $s^{-1}$) and greater variability, including slight sequence-dependent effects (**Figure 53a, c**). This trends suggest that $\tau_e$ is governed in general more strongly by amino acid type than by sequence context.

Analysis of $\Gamma_2$ profiles, which reflect the order parameter $S^2$, clearly highlights the heterogeneity in dynamic behaviour across all three amino acid types (**Figure 53b, d and f**), emphasizing the complexity of methyl side-chain motion in MALT1. This variability enables the use of $\Gamma_2$ as a benchmark for evaluating theoretically back-calculated parameters and comparing them across trajectories ensembles **4** and **14**.

Inspection of Figure **S3** indicates that, overall, trajectory **4** aligns closely with the experimental data, falling in general within the range of experimental uncertainties for both $R_1$and $\Gamma_2$. We consider this result important from a methodological point of view, as it indicates that free MD dynamics reproduces seemingly correctly the dynamics of methyl side chains for Val, Leu and Ile residues. This finding is particularly notable given that force field parameters for methyl groups are still under investigation. Moreover, the result expected to be if methyl dynamics in MALT1(PCASP-Ig3)$_{338–719}$ are consistent with the backbone dynamics observed in ensemble of trajectory **4.**

It is essential to emphasize that the other trajectories, corresponding to conformational ensemble **14**, obtained under higher salt conditions and exhibiting distinct active-site conformations but similar **W580** aromatic ring orientations, also align well with the experimental data (**Figure S5**). A striking example is the nearly identical $\mathbf{R_1}$ and $\mathbf{\Gamma_2}$ relaxation parameters for **Ile501** (**Figure5e, f**) across two ensembles (**4** and **14**), despite the substantial repositioning of this residue in the active versus inactive conformation. This suggests that the **Loop 3** conformational transition in the **PCASP** domain has minimal impact on the fast methyl group dynamics.

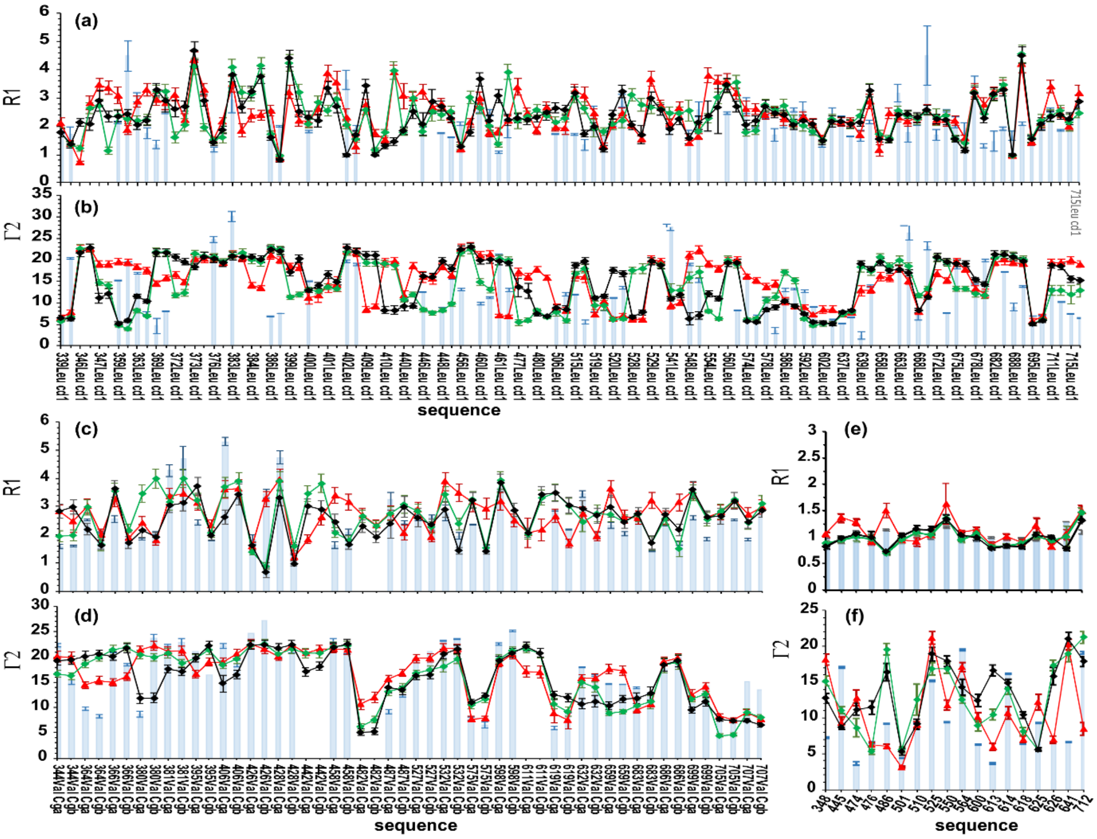


**Figure S5 MALT1(PCASP-Ig3)$_{339—719}$ Ile, Val; Leu Methyl Dynamic Parameters of the $R_1$ and $\Gamma_2$ obtained on 800 MHz spectrometers.** In Panels (a, c, and e) and (b, d and f) are presented experimental (light blue) and back calculated relaxation parameters $R_1$ and $\Gamma_2$, respectively of methyl Leu (a,b), Val (c, d) and Ile(e,f), extracted from trajectories 4 (red), 8(green) and 9(black).

### Relaxation Analysis Across Hydrophobic Clusters

To identify methyl groups most affected by interdomain interactions between **Ig3** and **PCASP**, we sorted **Val, Leu, and Ile** methyl's into four distinct hydrophobic clusters (**Cl1–Cl4**, **Figure S6**). These clusters are: **Cl1** (yellow) within the **Ig3** domain, **Cl2** (violet) at the **Ig3–PCASP** interface, and **Cl3** and **Cl4** (green and red, respectively) positioned on opposite sides of the **PCASP** β-sheet (**Figure S6**).

For **Cl1**, which is distant from the major conformational differences between ensembles **4** and **14**, the deviations between experimental and back-calculated $\mathbf{R_1}$ and $\mathbf{\Gamma_2}$ values are expected to be minimal (**Figure S6d1, d2**).

Clusters **Cl3** and **Cl4** (**Figures S6a1,a2 and b1, b2**), where experimental $\mathbf{R_1}$ and $\mathbf{\Gamma_2}$ data are limited, show minor variations among the back-calculated parameters for available methyl groups. This suggests that the **Loop 3** conformational changes exert little influence on fast dynamics within these clusters.

Similarly, for surface-exposed methyl groups (**Figure S6e1, e2**), both trajectory ensembles (**4** and **14**) exhibit strong agreement with experimental data, indicating that salt concentration does not significantly affect the fast dynamics of these methyls.

### Cl2: A Critical Cluster at the Ig3–PCASP Interface

The Cl2 cluster is particularly significant as it is located at the Ig3–PCASP interface, near the active site. Previous studies (Wallerstein *et al.*, 2024) suggest that Cl2 plays a crucial role in MALT1(PCASP-Ig3)$_{338–719}$ interactions with allosteric ligand. Nevertheless, the relaxation parameters $R_1$ and $\Gamma_2$ of the Cl2 methyl groups (**Figures S6c1, c2**) reveal that the back-calculated values for both trajectories (4 and 14) similar and closely match the experimental data.

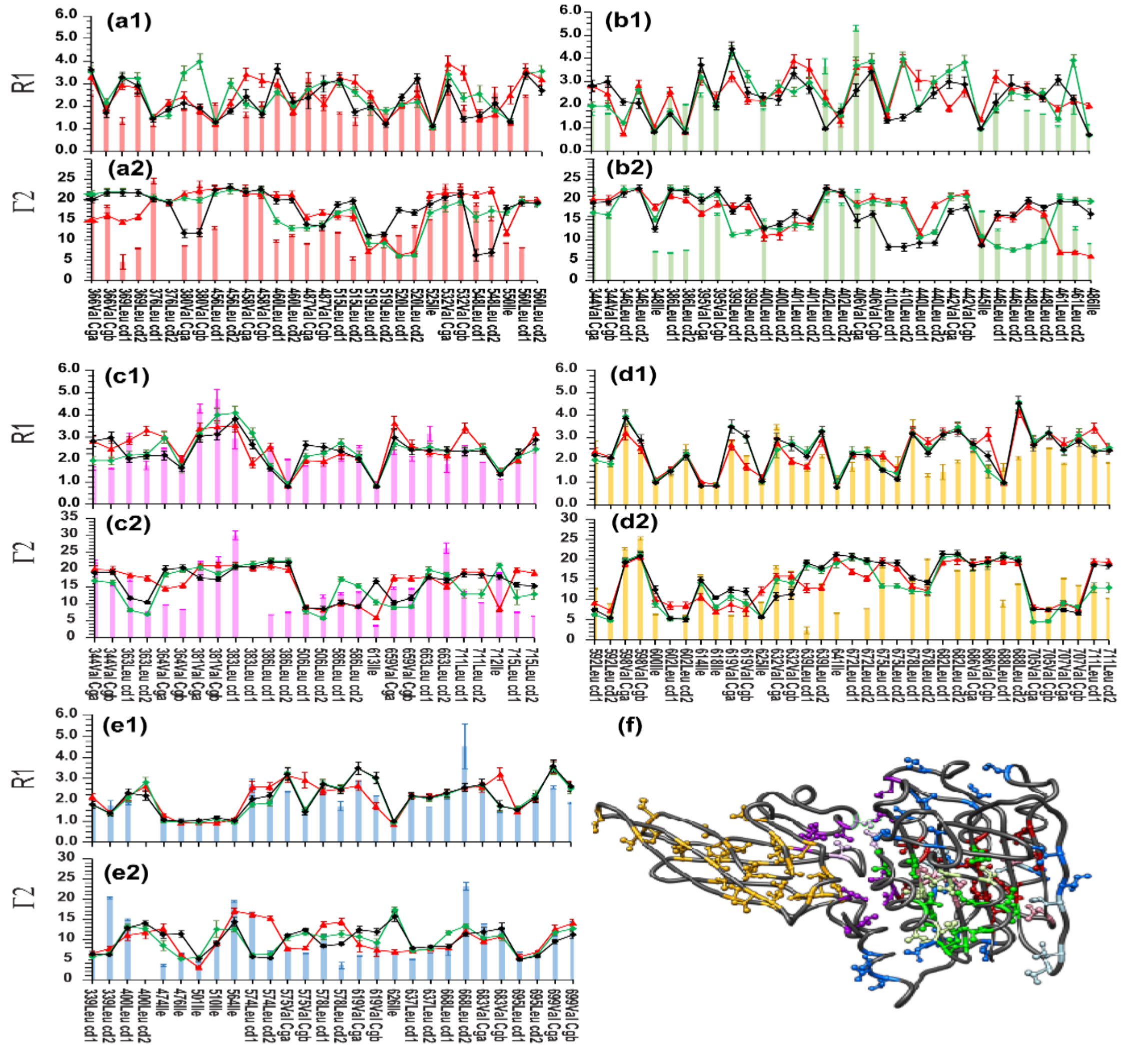


**Figure S6. Methyl relaxation dynamics across hydrophobic clusters in MALT1(PCASP-Ig3)$_{339–719}$** (a1–e1) Experimental $R_1$ relaxation rates (800 MHz, 25 °C) for methyl groups grouped by hydrophobic clusters: Cl3 (a1), Cl4 (b1), Cl2 (c1), Cl1 (d1), and surface-exposed methyl groups (e1). (a2–e2) Corresponding experimental $\Gamma_2$ relaxation rates. Experimental values are shown as colour-coded brackets matching the cluster colours in panel (f). Back-calculated relaxation parameters from MD trajectories are shown as solid lines: red (trajectory **4**), green (trajectory **8**), and black (trajectory **9**). (f) Spatial distribution of methyl clusters mapped onto the MALT1 structure. Cl1 (yellow) lies within the Ig3 domain; Cl2 (violet) occupies the Ig3–PCASP interface; Cl3 (green) and Cl4 (red) lie on opposite faces of the PCASP β-sheet. Surface-exposed methyl groups not participating in cluster formation are shown in blue. Assigned methyl groups are indicated with darker shades; unassigned methyl's with lighter shades. Error bars for experimental data represent one standard deviation from curve fitting; error bars for calculated parameters correspond to bootstrap-derived uncertainties.

### Assessment of the Oligomeric State of MALT1(PCASP-Ig3)$_{339\text{-}719}$ at High Salt Concentration

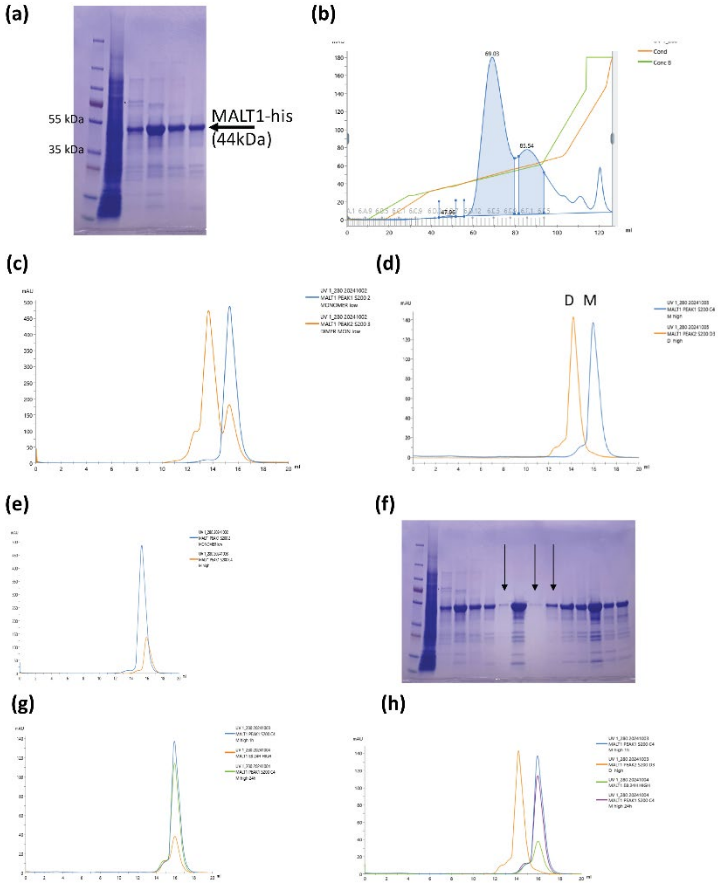


**Figure S7 Salt dependence of the monomeric state of the apo MALT1(PCASP-Ig3)$_{339\text{–}719}$ based on Ni-NTA affinity purification** (a,b) Gel and QXL column profiles of MALT1(PCASP-Ig3)$_{339}$ in buffer buffer: 20 mM Tris 7.5, 150 mM NaCl, 1 mM TCEP. Peak 1(D12-E7) corresponds to Monomer, Peak2 (E12-E5) corresponds to Dimer form of protein. (c,d) (D12-E7 of QXL column) and (E12-E5 of QXL column) were concentrated and loaded onto SEC200, shows monomer peak (blue) and dimer peak ( orange) in buffer (c) 20 mM Tris 7.5, **50 mM NaCl**, 1 mM TCEP and (d) 20 mM Tris 7.5, **500 mM NaCl**, 1 mM TCEP. MALT1 maintained as the same polymerization status in 50 and 500mM NaCl conditions. Profile of monomeric MALT1 form in different buffer as in (c,d) after 1 hour incubation (e) and 24 hour (g).

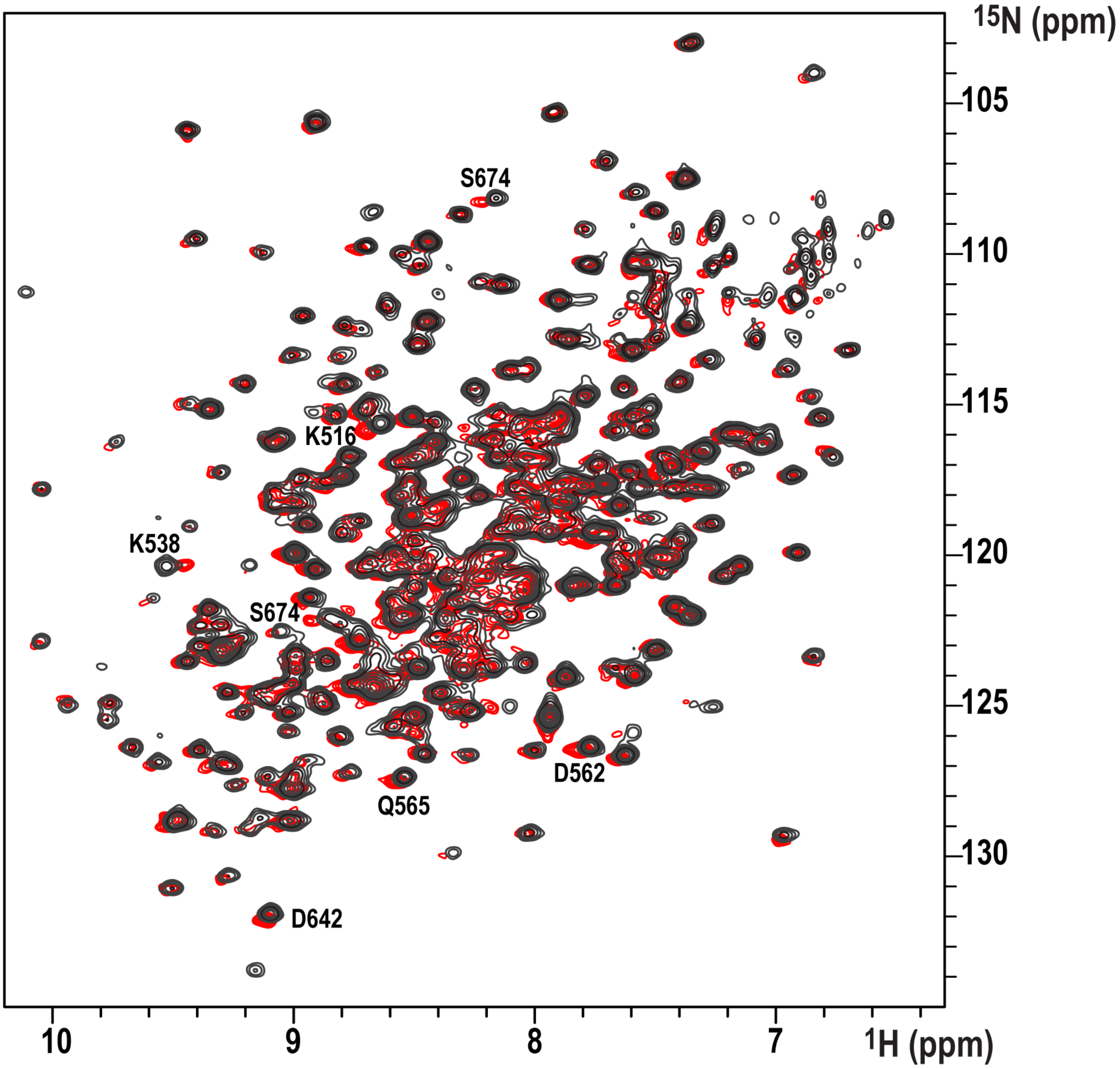


**Figure S8 | TROSY $^{1}H$–$^{15}N$ HSQC spectra of $^{15}N$-labeled MALT1 at different salt concentrations.** Spectra were recorded at low salt (black; 60 mM NaCl; 55 $Na^{+}$, 41 $Cl^{-}$) and high salt (red; 500 mM NaCl; 415 $Na^{+}$, 401 $Cl^{-}$) conditions. Amino acids exhibiting slight chemical shift perturbations are labeled.

### MD Simulation: Effect of Trajectory Length and Selection on the Back-Calculation of Relaxation Parameters for MALT1(PCASP-Ig3)$_{339–719}$

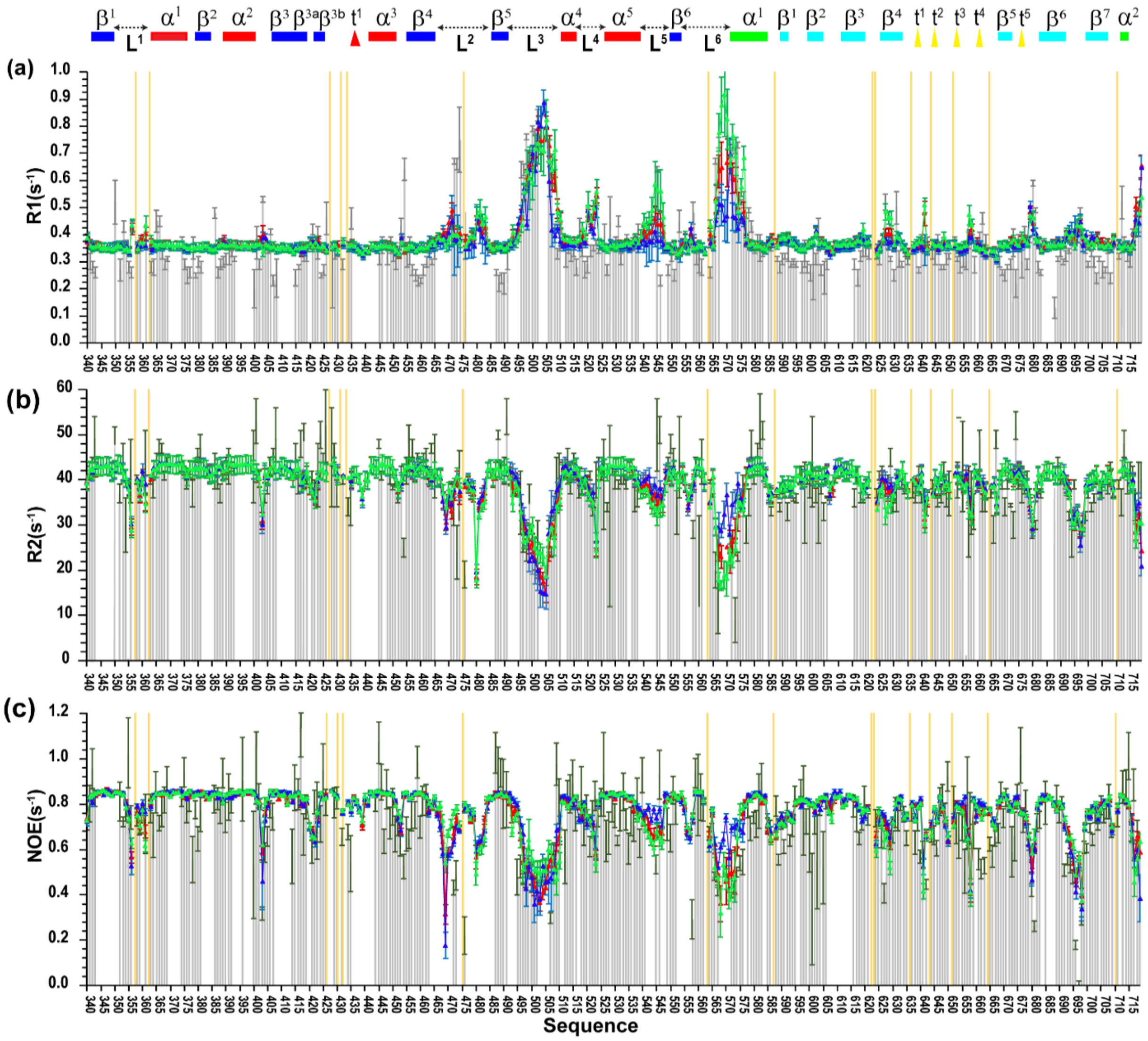

**Figure S9 MALT1(PCASP-Ig3)$_{339–719}$, amide backbone, $^{15}$N(H), dynamic parameters obtained on 800 MHz spectrometers.** The relaxation parameters for the MALT1(PCASP-Ig3)$_{339–719}$ are presented according to the following: longitudinal relaxation rate $R_1(s^{-1})$, the transverse relaxation time $R_2(s^{-1})$ and heteronuclear $^1H–^{15}N$ NOE values are presented in (a), (b) and (c), respectively. The experimentally obtained $R_1(s^{-1})$, $R_2(s^{-1})$ and NOE values are presented by light green solid brackets. The theoretically predicted dynamic parameters $R_1(s^{-1})$, $R_2(s^{-1})$ and NOE, obtained through three trajectories using starting structure for trajectory **4**, are displayed by solid lines and coloured in red, blue and black for ensembles obtained on segments of MD trajectories: 1000–1500 ns, 1000–3000 ns, and 2500–3000ns, respectively. The error bars of the experimental data are one σ from the curve fitting and for the predicted parameters from the bootstrapping analysis. Yellow bars show the position of proteins.

### Effect of Force Field

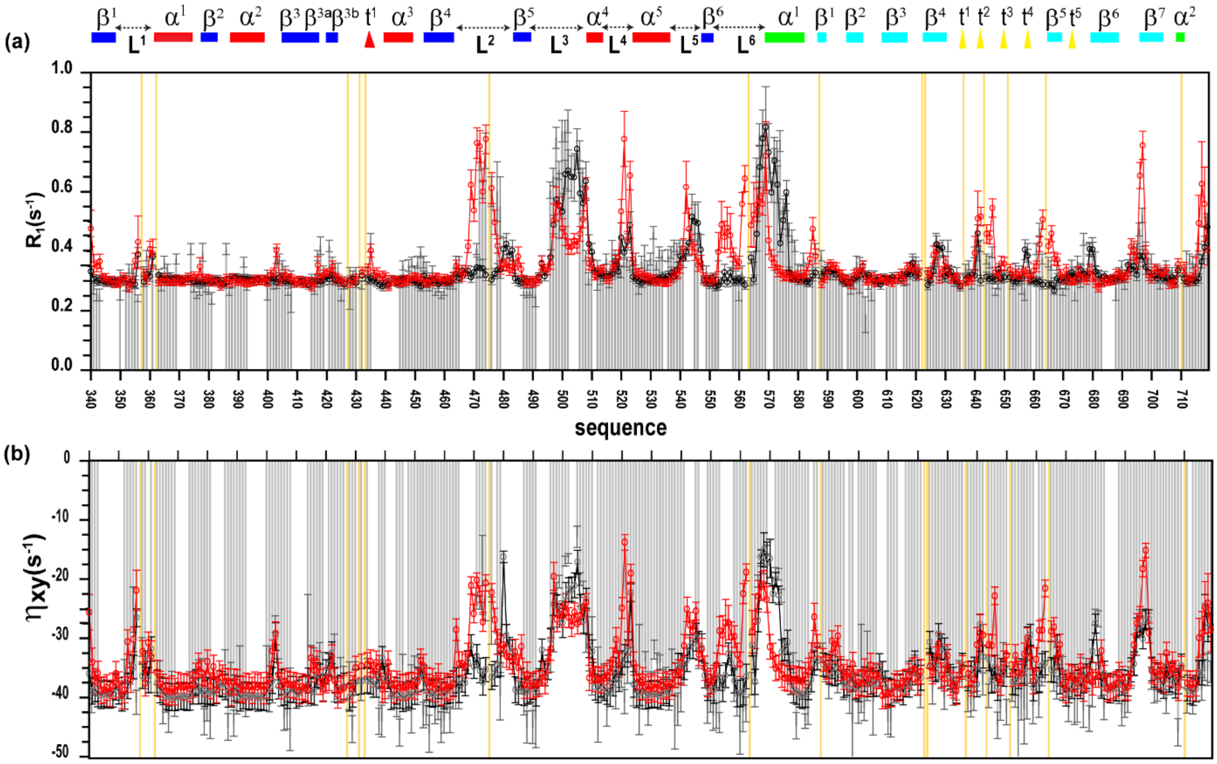


**Figure S10 MALT1(PCASP-Ig3)$_{339–719}$, amide backbone, $^{15}$N(H), dynamic parameters obtained on 900 MHz spectrometers**. The relaxation parameters for the MALT1(PCASP-Ig3)$_{339–719}$ are presented according to the following: longitudinal relaxation rate $R_1(s^{-1})$ and CSA/Dipole cross correlation relaxation, $\eta_{xy}$, values are presented in (**a**) and (**b**) respectively. The experimentally obtained $R_1(s^{-1})$ and $\eta_{xy}$ $(s^{-1})$ values are presented by light grey solid brackets. The theoretically predicted dynamic parameters $R_1(s^{-1})$ and $\eta_{xy}$ $(s^{-1})$, obtained through two **trajectories 4,** and **5**, are displayed by solid lines and coloured in black and red. The ensembles **4** obtained on segments of MD trajectories **2500–3000ns in force field CHARMM36**, and for ensemble **5** obtained on segments of MD trajectory **2500-3000ns in AMOEBA force field,** respectively. The error bars of the experimental data are one σ from the curve fitting and for the predicted parameters from the bootstrapping analysis.
Yellow bars show the position of proteins.

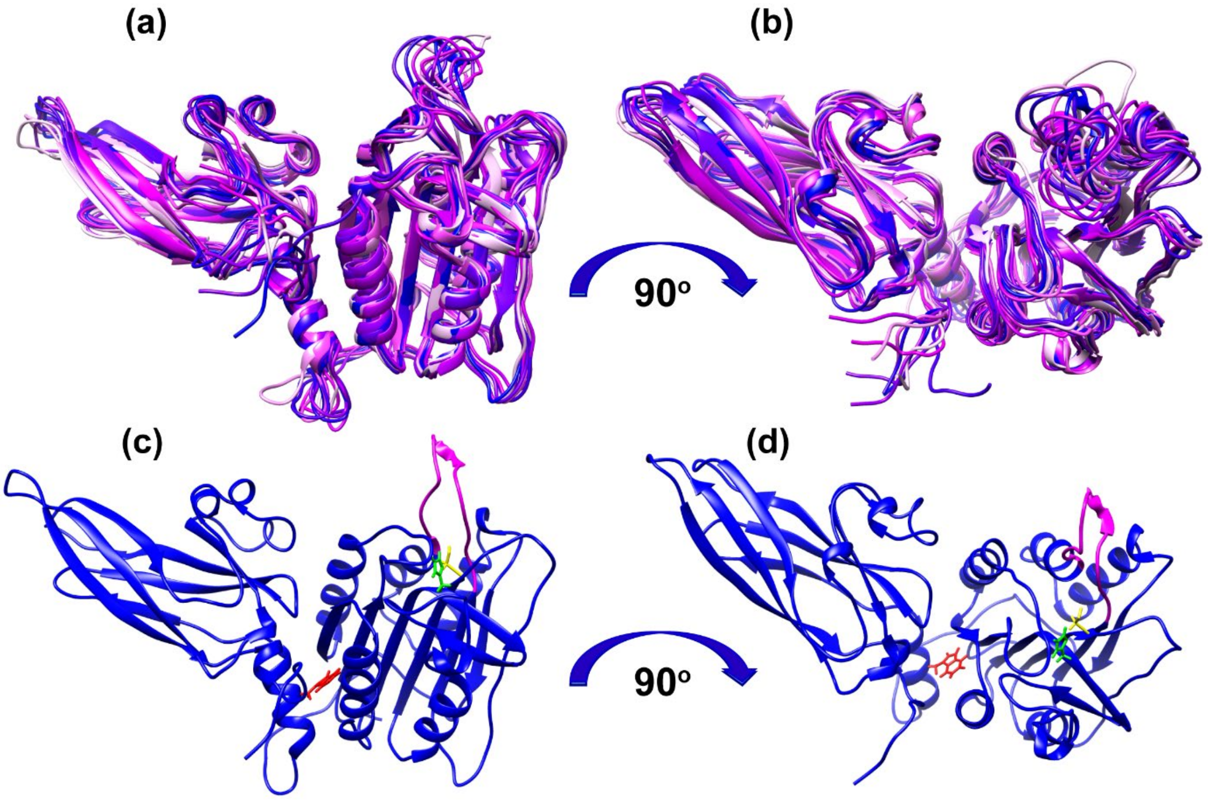


**Figure S11. Conformational ensembles of MALT1(PCASP-Ig3)$_{339–719}$ were obtained from free-restraints MD simulations at low salt concentration.** (**a**) Superposition of 10 structures from **trajectory 4**, identified through cluster analysis as the most populated ensembles. (**b**) The same structures as in (**a**) rotated by 90°. (**c**) Structure in the most populated ensemble from trajectory **4**, identified through cluster analysis. (**d**) The same structure as in (**c1**) rotated by 90°. In the structures shown in (c, d), the following amino acids are colored as follows: W580 in red, catalytic H415 in green, C464 in yellow, and loop 3 residues in pink. The **4**-th trajectory (2500-3000 ns time interval) was clustered into 20 states using an RMSD cutoff of 0.105 nm. The cluster populations were: 1–34.4%, 2–17.7%, 3–13.5%, 4–8.9%, 5–4.4%, 6–3.5%, 7–3.3%, 8–2.7%, 9–1.6%, 10–1.3%, 11–1.0%, 12–1.0%, 13–0.9%, 14–0.7%, 15–0.6%, 16–0.5%, 17–0.5%, 18–0.5%, 19–0.4%, 20–0.3%, and other–2.3%. The most representative 20 structures as a result of cluster analysis of the **4**-th trajectory (time interval 2500-3000 ns) have been deposited in the Protein Data Bank (PDB) (Entry ID: D_4-R5HP).
